# The PGC-1α Gly482Ser rs8192678 SNP reprograms oxidative metabolism to favor energy storage in females

**DOI:** 10.64898/2026.10.16.759626

**Authors:** Maria Galipeau, Nathalie Jouvet, Matthieu Schoumacher, Emilie Courty, Roxanne Vandenbeek, Aysim Gunes, Martin Braud, Cynthia Fourgeux, Jeremie Poschmann, Francis C. Lynn, May Faraj, Jennifer. L. Estall

## Abstract

The Gly482Ser SNP (rs8192678) in the peroxisome proliferator-activated receptor gamma coactivator 1 alpha (*PPARGC1A)* gene has been associated with obesity, type 2 diabetes and other human metabolic diseases. In this study, we found that human iPSCs carrying a serine risk allele had reduced respiratory capacity, and individuals living with overweight or obesity homozygous for the serine allele (S/S) exhibited a preference for carbohydrate oxidation following a high-fat meal. Using a PGC-1α^Ser481Gly^ knock-in mouse model, we performed metabolic phenotyping of male and female mice carrying the three genotypes (G/G, S/G, S/S) under acute nutritional challenges (chow, fasting and high-fat high-sugar (HFHS) feeding) and chronic HFHS feeding. S/S females were heavier and showed a trend toward an increased respiratory exchange ratio (RER) during fasting, with greater weight gain and fat mass following chronic HFHS feeding. Transcriptomic analyses revealed tissue-specific reprogramming of mitochondrial oxidative pathways, that were downregulated in subcutaneous adipose tissue and liver while upregulated in muscle, with the strongest effects observed in S/S females. Together, these findings provide *in vivo* evidence that this SNP reprograms whole-body oxidative metabolism, with a net effect of favoring carbohydrate oxidation and energy conservation, particularly in females.

**Highlights:**

- Human iPSC lines expressing the PGC-1α S482 variant had reduced respiratory activity.
- The PGC-1α Gly482Ser SNP was associated with an increased respiratory quotient in people living with overweight or obesity following a high-fat meal.
- This PGC-1α SNP exerted sexually dimorphic effects on energy metabolism, with female PGC-1α^Ser481^ mice displaying impaired metabolic adaptation to nutritional challenges, including reduced fasting lipid oxidation and lower energy expenditure during HFHS refeeding.
- Transcriptomic analysis revealed multi-tissue reprogramming of mitochondrial and fatty acid beta-oxidation pathways in PGC-1α^Ser481Gly^ mice.
- The PGC-1α S481 variant may predispose females toward energy conservation, providing a physiological basis for association of this variant with increased metabolic disease risk.

## Introduction

Obesity and type 2 diabetes (T2D) are multifactorial, polygenic and interconnected chronic diseases. Evidence from twin studies suggests that genetic factors explain 30–90% of interindividual differences in disease susceptibility (1–3), yet variants identified by genome-wide association studies explain only a fraction of estimated heritability. The remainder may reflect variants whose metabolic consequences only emerge under specific environmental conditions, such as dietary excess, physical inactivity, or aging (4–7). This context-dependence underscores the importance of gene-environment interactions in shaping metabolic disease risk, with sex further influencing disease etiology, evidenced by higher prevalence of obesity in females and T2D in males (8, 9). A better understanding of how genetic variants interact with diet, lifestyle, and sex to alter metabolism is essential to develop personalized strategies for genetically predisposed individuals.

A compelling example of such genetic variance is the PGC-1α Gly482Ser SNP (rs8192678, C>T or G>A), whose contribution to metabolic disease risk appears to be influenced by physical activity, age, and sex across independent clinical studies (10–14). This common missense SNP encodes either a glycine (Gly; G) or a serine (Ser; S) amino acid at position 482 of the peroxisome proliferator-activated receptor gamma coactivator 1 alpha (PGC-1α) protein (Fig. 1a), a transcriptional coactivator that coordinates expression of metabolic gene programs to maintain energy homeostasis, including mitochondrial biogenesis, oxidative phosphorylation, and macronutrient oxidation (15–18). The serine (S482) risk allele, present at a global frequency of approximately 30% (ALFA project 2026 n=623 668 (19)), is associated with T2D susceptibility, weight gain, insulin resistance, and obesity across multiple ethnicities (20–22, 13, 23, 24). The S482 allele is also linked to decreased PGC-1α mRNA in human pancreatic islets, liver and skeletal muscle (12, 25, 26), though whether reduced PGC-1α expression represents a cause or a consequence of metabolic dysfunction remains to be determined.

**Figure 1.**
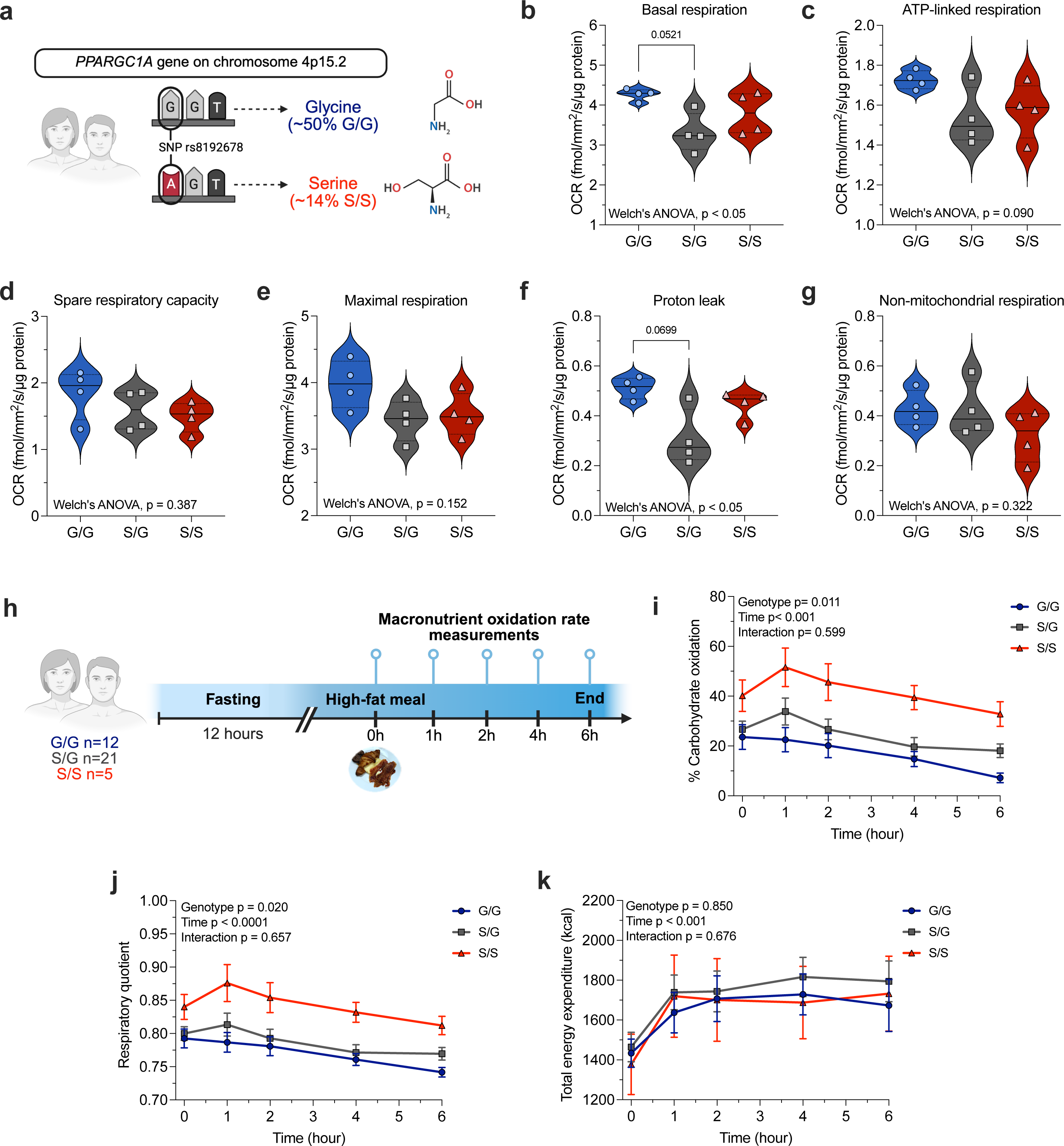
The PGC-1α Gly482Ser SNP is associated with reduced cellular respiration and higher postprandial carbohydrate oxidation in humans. **a)** Codon change introduced by the PGC-1α Gly482Ser SNP rs8192678 and corresponding amino acid two-dimensional structures. The GGT (glycine) to AGT (serine) substitution shown reflects the coding (minus-sense) strand. Global frequency (%) of homozygous genotypes were estimated from ALFA allele frequencies using Hardy-Weinberg equilibrium (96, 97). **b**) Basal respiration, **c**) ATP-linked respiration, **d**) spare respiratory capacity, **e**) maximal respiration, f) proton leak and g) non-mitochondrial respiration in G/G, S/G and S/S human iPSCs (n=4 independent clones per genotype). Respiration parameters were derived from OCR measurements and normalized to total cellular protein content. Values for each clone were averaged across 3-4 independent experiments. **h**) Schematic of the human study design. Following a 12-h fast, participants consumed a high-fat meal and macronutrient oxidation was measured over 6 h by indirect calorimetry. **i**) Carbohydrate oxidation, **j**) Respiratory quotient and **k**) Energy expenditure during the 6-h postprandial period. Data are shown as mean ± SEM. Statistics: **b-g**) Welch’s ANOVA with Dunnett T3 multiple comparisons test; **i-k**) Two-way repeated measures ANOVA with Bonferroni’s multiple comparisons test.

Under physiological conditions, PGC-1α expression is very dynamic, induced by fasting, exercise, cold, and acute inflammation, and suppressed upon refeeding, aging and chronic inflammation (15, 27, 28, 12, 29–32). The protein is also quickly degraded and we have previously shown that the S482 variant reduces PGC-1α protein stability *in vitro* (33), consistent with reduced coactivation of *Pepck*, *Cpt1*, *Sod2*, and *Gpx1* (34, 33, 35). Independently*, PPARGC1A* expression is reduced in subcutaneous adipose tissue, pancreatic islets, and skeletal muscle of individuals living with obesity and T2D (36, 37, 25), further supporting an association between reduced PGC-1α expression and metabolic disease. Conversely, other groups have shown that the S482 variant increases PGC-1α protein and mRNA levels, promoting lipogenesis and mitochondrial respiratory capacity in adipocytes (38), and fat accumulation in hepatocytes (34). Together, these findings support that the PGC-1α S482 variant meaningfully alters PGC-1α expression and function in different cell types. However, the downstream physiological and biological consequences of this variant across tissues, and how it translates into changes in whole-body energy metabolism and metabolic disease susceptibility, remain poorly understood.

In this study, we found that the PGC-1α S482 variant was associated with reduced cellular respiration and lipid oxidation in mice and humans. Data in mice revealed tissue-specific transcriptional reprogramming of mitochondrial oxidative metabolism linked to the SNP, and that the metabolic phenotype associated with the serine allele likely supports lipid conservation or storage under conditions of dietary caloric excess. Together, our work provides the first *in vivo* characterization of energy metabolism associated with the PGC-1α Gly482Ser SNP and offers mechanistic insight into its contribution to metabolic disease risk.

## Results

### Human iPSCs carrying the serine allele have reduced respiratory activity

Given the central role of PGC-1α in regulating energy metabolism and mitochondrial function, we assessed the impact of the PGC-1α Gly482Ser SNP on oxygen consumption rate (OCR) in isogenic G/G, S/G and S/S clones engineered from a single male human iPSC line (originally S/G genotype). Four independent, undifferentiated clones per genotype were analyzed using a continuous metabolic monitoring system (Resipher). S/G iPSCs exhibited lower basal respiration and proton leak compared to G/G iPSCs, with trends toward reduced ATP-linked and maximal respiration. However, spare respiratory capacity and non-mitochondrial respiration were not significantly affected by genotype (Fig. 1b-g). Because many clinical studies have identified phenotypes associated with the serine allele using a dominant genetic model (G/G vs S/G *+* S/S) (39–41, 24, 42), we also assessed OCR-related parameters after pooling serine variant expressing clones (S/G and S/S) (Fig. S1). Under this genetic model, iPSCs carrying at least one serine allele exhibited significantly lower basal respiration, ATP-linked respiration and proton leak compared to G/G iPSCs, with a trend toward lower maximal respiration (Fig. S1a-e). Spare respiratory capacity and non-mitochondrial respiration remained comparable between groups (Fig. S1c,f). Together, these results demonstrate that the serine allele is associated with reduced cellular respiration in human pluripotent cells.

### S/S individuals living with overweight or obesity oxidize less lipid after a high-fat meal

Reductions in cellular respiratory capacity can influence how the body metabolizes and stores macronutrients (43, 44). Decreases in respiration can cause a switch toward anaerobic metabolism and increase reliance on carbohydrate fuels, since beta-oxidation requires an efficient electron transport chain (45, 46). To test whether the decrease in respiratory activity linked to the S482 variant is associated with changes in whole-body macronutrient metabolism, we ran post-hoc analysis of baseline data and DNA samples from people living with overweight or obesity (without cardiometabolic disease) who participated in two clinical trials at the IRCM, as published (47, 48). Genotyping of these research participants yielded the three genotype groups with the expected incidence ratio (G/G n=12, S/G n=21, S/S n=5) within the Hardy-Weinberg equilibrium (χ^2^ = 0.789, p=0.674). Anthropometric and fasting metabolic parameters did not differ significantly across genotypes (Table 1).

**Table 1.**
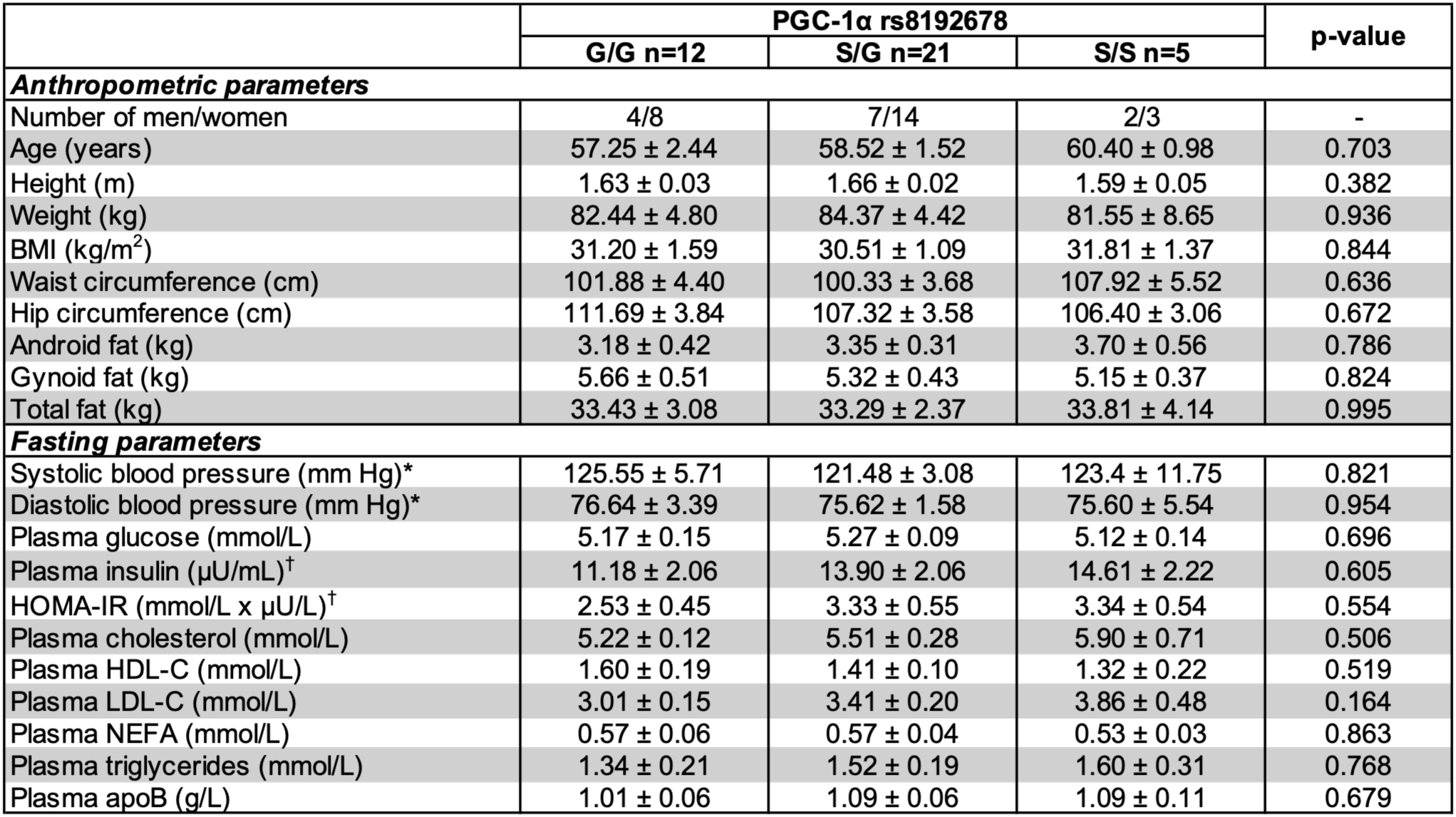
Anthropometric and fasting metabolic parameters were similar between G/G, S/G and S/S individuals. Data are presented as mean ± SEM. Statistics: One-way ANOVA; p ≤ 0.05 was considered statistically significant. For missing data: * for n = 11 G/G and † for n = 20 S/G. Abbreviations: BMI: body mass index; HOMA-IR: homeostatic model assessment of insulin resistance; HDL-C: high-density lipoprotein cholesterol; LDL-C: low-density lipoprotein cholesterol; NEFA: non-esterified fatty acids; apoB: apolipoprotein beta.

Following a 12-hour fast, participants consumed a standardized high-fat meal and macronutrient oxidation rates were measured over 6 hours by indirect calorimetry (Fig. 1h). S/S carriers had significantly higher carbohydrate oxidation, and consistently, a higher respiratory quotient following a high-fat meal compared to S/G and G/G carriers (Fig. 1i,j), which remained significant after adjustment for BMI, age or sex. There was no significant difference in total energy expenditure (Fig. 1k). Taken together, these findings suggest that the metabolic consequences of the serine allele extend from the cellular to the organismal level. Although derived from distinct experimental systems, the reduced respiratory activity observed in iPSCs together with altered substrate utilization observed in human carriers, reveal important functional consequences of this SNP on PGC-1α mediated control of energy metabolism.

### S/S female mice oxidize less lipid during fasting and are prone to energy conservation when refed a HFHS diet

To bridge our cellular and human physiological findings, we generated a PGC-1α^Ser481Gly^ knock-in mouse model. Because the murine *Ppargc1*α gene naturally encodes a serine residue at the orthologous position (S481), our PGC-1α^Ser481Gly^ mouse model is uniquely suited to directly test whether substituting a glycine residue at this position (in one or both alleles) affects PGC-1α expression, nutrient metabolism and/or metabolic outcomes. Given our previous findings that overexpressed PGC-1α S482 variant has a decreased protein half-life (33), we expected the PGC-1α^Ser481Gly^ knock-in mice to have reduced PGC-1α expression, with a similar but subtler phenotype than mice with complete PGC-1α knockout (49, 50). Unexpectedly, only female S/G and S/S mice showed blunted glucagon-induced PGC-1α protein levels in hepatocytes compared to G/G female mice (Fig. 2a,b), while there were no significant differences in PGC-1α protein levels in hepatocytes from male mice (Fig. 2c,d). These data suggest that differential regulation of PGC-1α expression could result in sex-specific metabolic consequences of the SNP.

**Figure 2.**
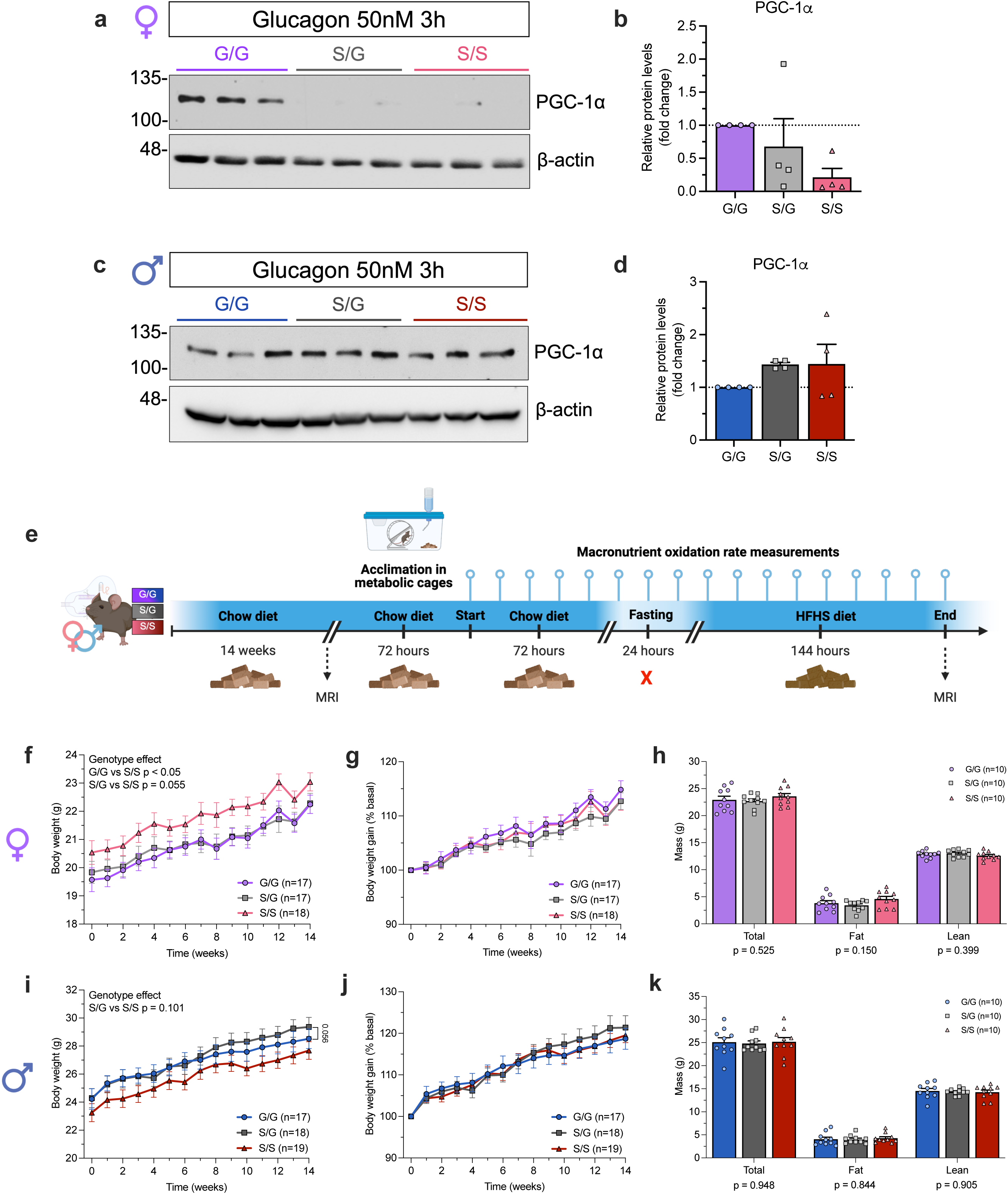
Female S/S mice are heavier during chow feeding despite similar body composition. Primary hepatocytes were isolated from 12-to 17-week-old female and male PGC-1α^Ser481Gly^ mice and treated with 50 nM glucagon for 3 h. Representative Western blots of PGC-1α protein levels in **a**) female and **c**) male hepatocytes (n=1 mouse per genotype in technical triplicates). Quantification of relative PGC-1α protein levels in **b**) female and **d**) male hepatocytes. Each dot represents the mean levels of an experiment (n = 4 biological replicates), each performed in technical duplicate or triplicate using hepatocytes pooled from 1-2 mice per genotype. **e**) Schematic of the mouse experimental protocol after 14 weeks of chow diet. Following 72 h of acclimation in Promethion metabolic cages, measurements were recorded for 72 h on chow diet, followed by a 24-h fast and 144 h of high-fat high-sugar (HFHS) refeeding. Weekly body weight (g) and body weight gain (%) of **f-g**) female and **i-j**) male PGC-1α^Ser481Gly^ mice over 14 weeks of chow feeding. Total body mass, fat mass, and lean mass (g) before acclimation in **h**) female and **k**) male mice. Statistics: **f,g,i,j**) Two-way repeated measures ANOVA; **h,k**) Genotype effect from one-way ANOVA, performed independently for each body composition parameter. Data are shown as mean ± SEM.

To assess the physiological and pathological consequences of the genetic variant on whole-body macronutrient metabolism, male and female PGC-1α^Ser481Gly^ mice were maintained on a chow diet for 14 weeks, after which indirect calorimetry was performed across three successive nutritional states: 72-hour *ad libitum* chow feeding, a 24-hour fast, and 144-hour *ad libitum* HFHS refeeding (Fig. 2e). Throughout chow feeding, S/S females had greater body weight compared to S/G and G/G mice (Fig. 2f), although this did not stem from differences in the trajectory of weight gain over time (Fig. 2g). Body composition prior to indirect calorimetry was similar across genotypes (Fig. 2h). In contrast, S/S males tended to have lower body weights compared to S/G males (Fig. 2i), again with comparable weight gain curves and body composition across genotypes (Fig. 2j,k). Together, these results point to a sexually dimorphic effect of the serine allele on body weight under basal conditions, with S/S females trending heavier and S/S males trending lighter.

To test whether S/S mice recapitulate the increased carbohydrate oxidation observed in human S/S carriers, we dynamically assessed the respiratory exchange ratio (RER) across successive chow, fasting and HFHS refeeding (Fig. 3a), conditions that challenge metabolic flexibility and reveal acute shifts in substrate utilization. RER was similar between female genotypes during chow feeding throughout the light and dark phases (Fig. 3b). However, upon fasting, S/S females showed a trend (p=0.175) toward higher RER across the entire fasting period (Fig. 3c). During HFHS refeeding, there were no significant differences in RER (Fig. 3d). Together, these findings support a role for the serine allele in regulating whole-body fuel substrate oxidation during periods of macronutrient deprivation.

**Figure 3.**
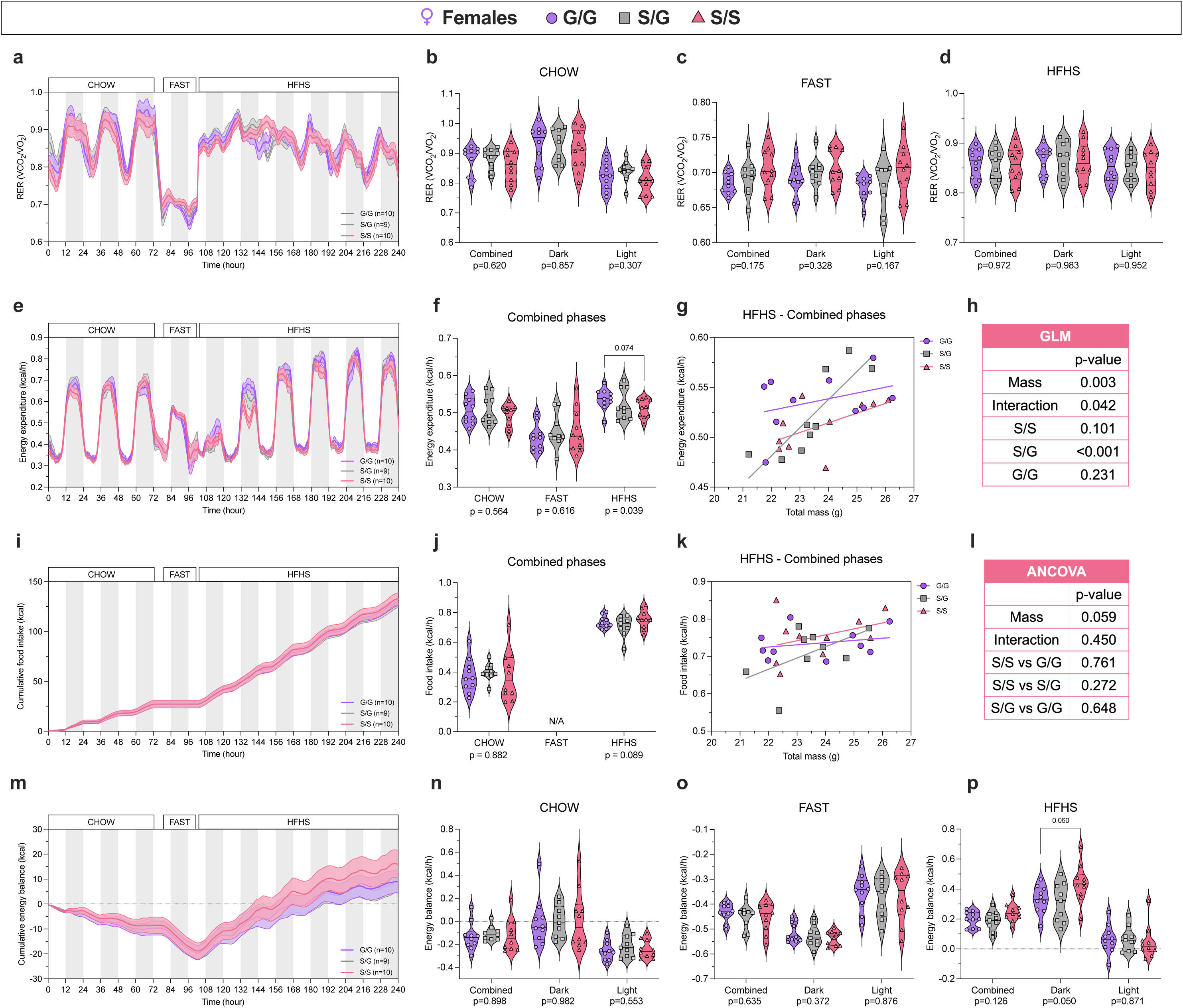
Lipid oxidation is lower in female S/S mice during fasting and is associated with positive energy balance under HFHS refeeding. **a**) RER versus time in female PGC-1α^Ser481Gly^ mice and violin plots representing the mean RER per mouse during **b**) chow feeding, **c**) fasting and **d**) HFHS refeeding per photoperiod. **e**) Energy expenditure (kcal/h) versus time. Violin plots representing the mean rates of **f**) energy expenditure per mouse during chow, fasting and HFHS combined photoperiods. **g**) Linear regression of mean energy expenditure rates against total body mass during the HFHS combined photoperiod and **h**) generalized linear model (GLM) of energy expenditure. **i**) Cumulative food caloric intake (kcal) versus time. Violin plots representing the mean rates of **j**) food intake (kcal/h) per mouse during chow, fasting, and HFHS combined photoperiods. **k**) Linear regression of mean food intake rates against total body mass during the HFHS combined photoperiod and **l**) ANCOVA of food intake. **m**) Cumulative energy balance (kcal) versus time. Violin plots representing the mean energy balance rates (kcal/h) per mouse during **n**) chow, **o**) fasting, and **p**) HFHS feeding per photoperiod. Shaded areas represent dark phases. Statistics: Genotype effects were assessed using **b**) Kruskal-Wallis test, **c,d,n-p**) one-way ANOVA with Tukey’s post hoc test, or **f,j**) ANCOVA with total body mass as a covariate, performed independently for each photoperiod or nutritional state.

Energy expenditure was similar between female genotypes during chow feeding and fasting (Fig. 3e,f). However, during HFHS refeeding, energy expenditure was reduced in S/S females compared to G/G females (Fig. 3f), and we detected a significant mass x genotype interaction (Fig. 3g,h). Using a generalized linear model (GLM), S/G females showed a significant positive relationship between body mass and energy expenditure (p < 0.001), whereas S/S and G/G females showed no significant mass-dependent relationship (p = 0.101 and p = 0.231, respectively) (Fig. 3h). Together, these results suggest that this PGC-1α SNP impacts energy expenditure during a HFHS challenge, and the blunted energetic response to macronutrient availability in S/S females may contribute to a positive energy balance under conditions of caloric excess.

Interestingly, while female mice of all genotypes ate similar amounts of food during chow feeding (Fig. 3i,j); HFHS food intake tended to be higher in S/S females even when adjusted for body mass (p = 0.089) (Fig. 3j). There was no significant mass x genotype interaction (p = 0.450) (Fig. 3k,l). Since we observed an influence of genotype on both energy expenditure and food intake, we examined whether this affected energy balance. Energy balance, defined as the difference between caloric food intake and energy expenditure, was similar during chow and fasting (Fig. 3m-o); however, S/S females on HFHS food showed higher energy balance during the dark phase compared to G/G females (Fig. 3p), suggesting a shift toward energy conservation when dietary fat and sugar are abundant. In contrast, S/S males showed no differences in RER, energy expenditure, food intake or energy balance across nutritional states (Fig. S2). Together, these results suggest a more energy-conserving metabolic phenotype in S/S females, characterized by trends toward higher food intake and lower energy expenditure in response to the switch to HFHS food. This phenotype was not observed in S/S males.

Since physical activity can significantly impact respiration, energy expenditure and food intake, we also assessed cage movement and voluntary wheel running over the calorimetry period (Fig. S3). During chow and HFHS feeding, all female genotypes travelled similar distances (and at similar rates) within the cage and in the wheel. Intriguingly, S/S females tended to have greater voluntary wheel running during fasting (Fig. S3a-h). Similarly, S/S males had higher cage movement during fasting and tended to run more in the wheel on chow (Fig. S3i-p). Together, these data suggest that S/S mice of both sexes have increased voluntary physical activity, particularly under conditions of nutrient limitation. Thus, energy expenditure was reduced in S/S females and unchanged in S/S males despite having increased physical activity, further supporting an energy conserving phenotype for the serine allele.

### RNA-seq reveals tissue-specific transcriptional reprogramming of mitochondrial and fatty acid oxidative pathways in PGC-1α^Ser481Gly^ mice under chow feeding

To gain insight into the mechanisms underlying metabolic differences in PGC-1α^Ser481Gly^ mice, we performed RNA sequencing of gastrocnemius muscle, subcutaneous adipose tissue (SCAT) and liver. These three tissues were selected for their central roles in lipid oxidation, storage and mobilization, and whole-body energy homeostasis (51). Male and female PGC-1α^Ser481Gly^ mice were maintained on chow diet for 24 weeks and fasted for 3 hours before tissue collection. Data from 3′ DGE-seq revealed modest sex differences in *Ppargc1*α mRNA expression across tissues, which were influenced by genotype in muscle and SCAT (Fig. S4a-c). Principal component analysis (PCA) revealed that transcriptional profiles clustered by sex along PC1 in muscle and liver; while in SCAT, sex separation was observed along PC2 and there was no clear genotype-dependent clustering in any tissue (Fig. S4d-f). Differential expression analysis comparing S/S and G/G groups in each tissue (FDR-adjusted p-values, Fig. S4g-i and Supplementary File 1) showed only one significantly differentially expressed gene in SCAT (Fig. S4h), *Tirap/Mal*, an adaptor protein linking Toll-like receptors to MyD88 signaling in innate immunity (52). The lack of strong gene changes in these samples was expected, given the small sample size, absence of a metabolic challenge, and the fact that this SNP does not result in major gain-or loss-of PGC-1α function (53).

To identify biological pathways that could be linked to genotype, we performed fast gene set enrichment analysis (fgsea) using the entire ranked gene list based on Wald statistics for each tissue. We identified 34 pathways commonly enriched across all three tissues in S/S vs G/G mice (Fig. 4a and Supplementary File 2). Among these, pathways related to mitochondrial oxidative metabolism (including the TCA cycle, electron transport chain and oxidative phosphorylation) were upregulated in muscle, but downregulated in SCAT and liver of S/S mice compared to G/G mice (Fig. 4b). Among the 58 pathways commonly enriched between muscle and SCAT, fatty acid beta-oxidation (FAβO) pathways were also upregulated in muscle but downregulated in SCAT (Supplementary File 2). Beyond bioenergetic alterations, the SCAT of S/S mice had increased inflammatory, immune, and cellular stress signaling pathways, specifically, enrichment of p53 signaling, MAPK signaling, mast cell activation, leukocyte migration, and leukocyte proliferation (Fig. 4b and Supplementary File 2). In liver of S/S mice, data suggested activation of intracellular stress cascades and structural modification programs, with upregulation of pathways involved in MAPK and JNK signaling, prostaglandin secretion, icosanoid transport, lipid export, blood vessel morphogenesis, and growth vasculature development (Fig. 4b and Supplementary File 2). Together, these data suggest that the PGC-1α serine variant coordinates a multi-tissue transcriptomic shift, characterized by overall downregulation of mitochondrial bioenergetics in liver and SCAT accompanied by increases in stress signaling, lipid export, and inflammatory pathways, contrasted by upregulation of oxidative metabolism and lipid catabolism genes in muscle.

**Figure 4.**
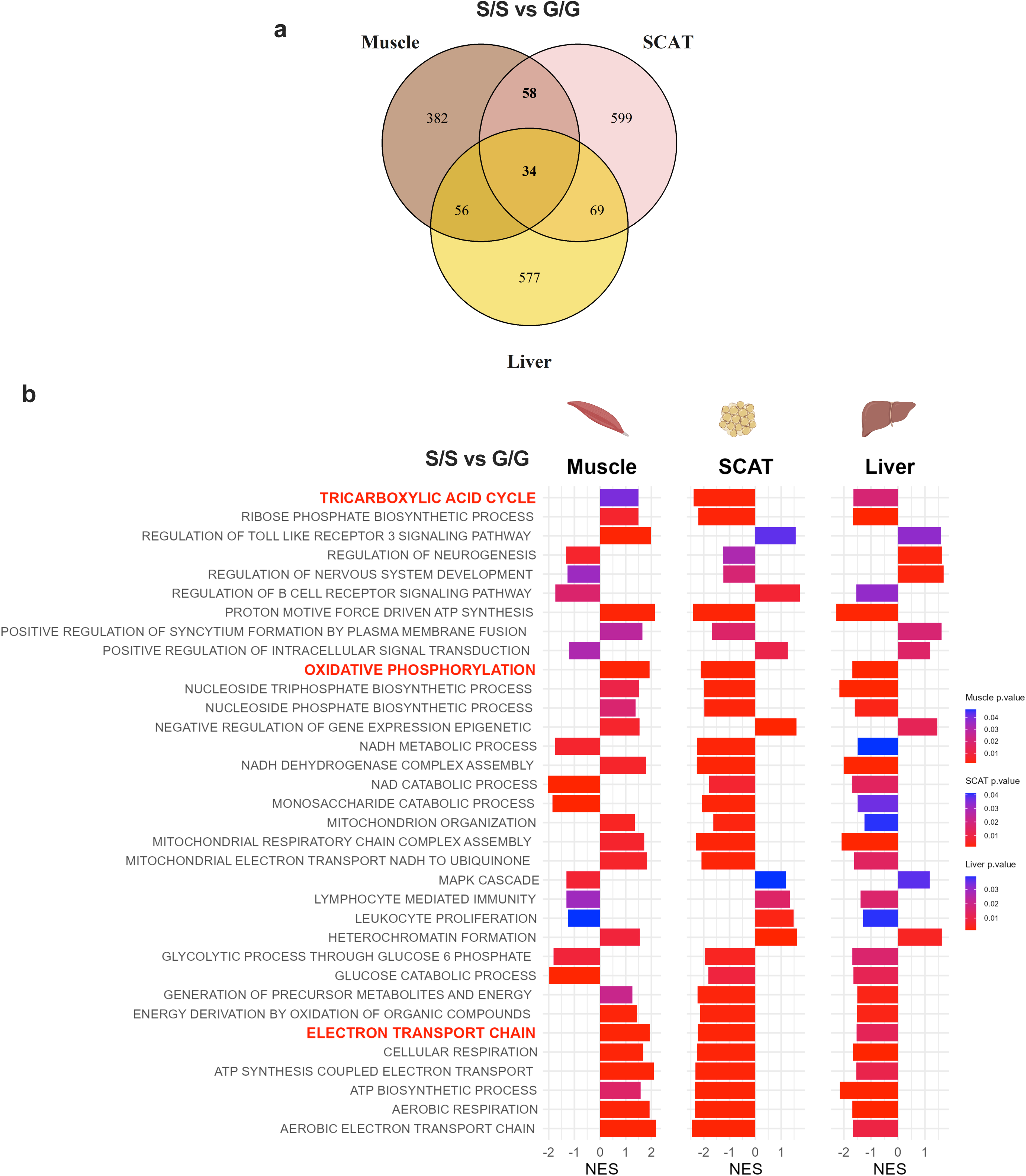
The PGC-1α S481 variant exerts opposing effects on mitochondrial oxidative pathways in liver, SCAT and muscle. **a**) Venn diagrams showing the overlap of differentially enriched pathways between G/G and S/S mice across gastrocnemius muscle, SCAT and liver tissues from PGC-1α^Ser481Gly^ mice, identified by fgsea using the GOBP gene sets (MSigDB M5 collection, 2024). **b**) Bar plot showing the 34 pathways commonly enriched between G/G and S/S mice across all three tissues. Bar length represents the normalized enrichment score (NES) and color indicates the raw p-value. n = 9-10 G/G, 9-10 S/G, 8-9 S/S mice.

Importantly, although we found sex to be a determinant in the physiological outcomes linked to the SNP, we did not stratify the above analysis by sex as it would have significantly reduced power. However, our data suggested that pathway regulation may be different between sexes. We and others have previously shown that sex influences the expression and function of PGC-1α (54, 33, 55, 56). Thus, we performed a targeted sub analysis to determine whether the pathways identified above behaved similarly in both sexes. For this, we generated heatmaps of genes involved in the tricarboxylic acid cycle, electron transport chain, oxidative phosphorylation and FAβO in each tissue (Fig. 5), where samples were grouped by genotype and sex. In muscle, both S/S male and female mice showed higher expression of genes involved in cellular respiration and FAβO compared to G/G mice (Fig. 5a). In SCAT, these pathways were downregulated in S/S of both sexes, but more strikingly in females (Fig. 5b). In liver, transcriptional profiles segregated primarily by sex, with females showing lower gene expression for electron transport chain and oxidative phosphorylation pathways, which was accentuated in serine allele carrying mice (Fig. 5c). While this analysis was limited to shared pathways across the tissues, we also noted that estrogen receptor signaling was one of the top regulated pathways in SCAT (raw p = 0.001, adjusted p= 0.127, Supplementary File 3). Using single-sample gene set enrichment analysis (ssGSEA), S/S females exhibited higher estrogen receptor signaling activity compared to G/G females in SCAT (Fig. S5b). As expected, estrogen receptor signaling activity was higher in female mice across all genotypes and tissues (Fig. S5a-c). Collectively, these observations suggest that the consequences of the SNP at the transcriptional level are similar between the sexes, whereas physiological differences, including reduced lipid oxidation and energy conservation phenotypes in females, may stem from hormonal influences on metabolic pathways.

**Figure 5.**
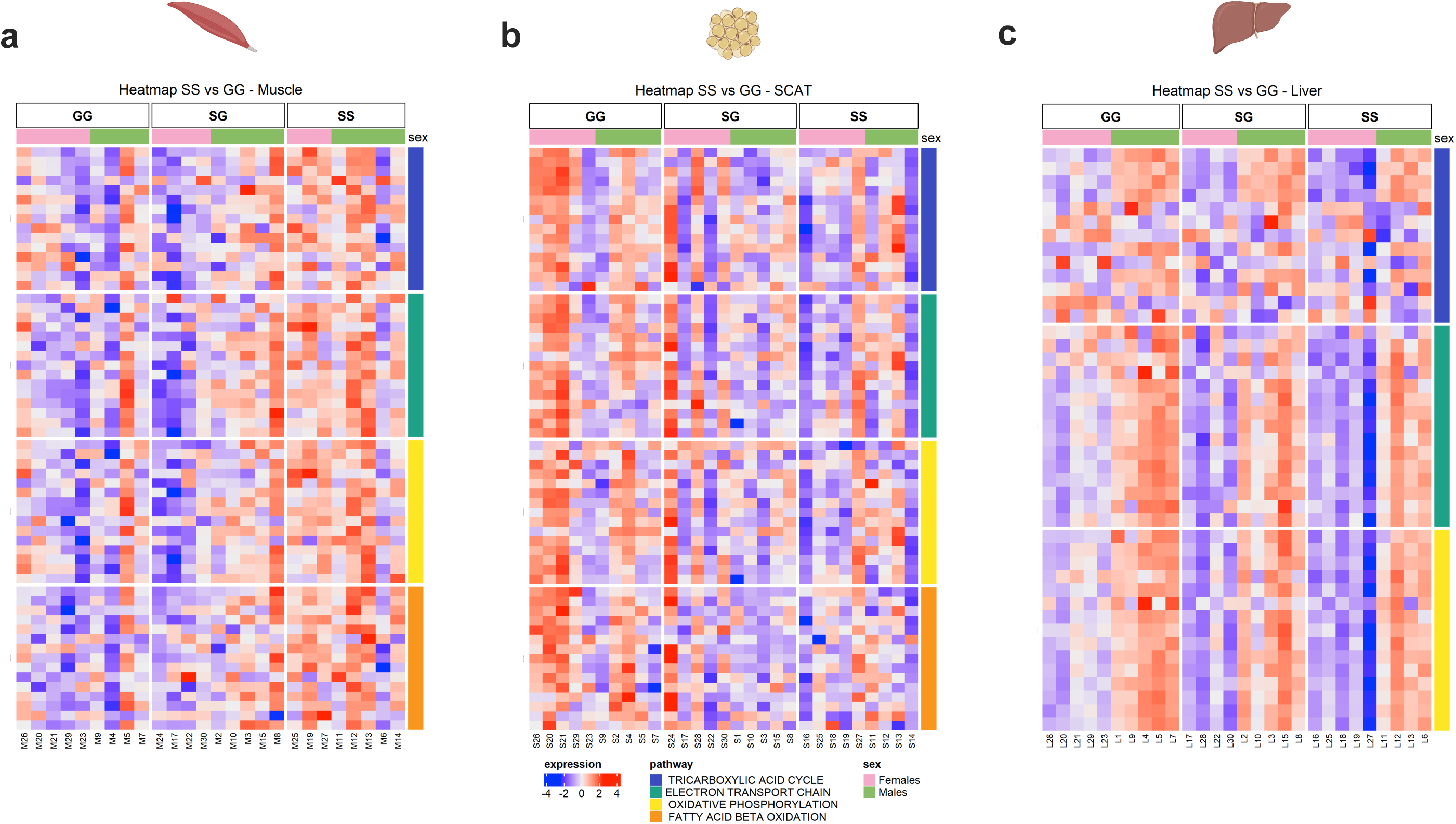
Heatmaps reveal opposing transcriptional effects on oxidative metabolism in muscle and SCAT in a sex- and genotype-dependent manner. Heatmap representation of the 15 most contributive genes (according to Wald statistics) to tricarboxylic acid cycle, electron transport chain, oxidative phosphorylation and fatty acid beta-oxidation pathways in **a**) muscle, **b**) SCAT and **c**) liver of PGC-1α^Ser481Gly^ mice of both sexes. Each row represents an individual gene and each column represents an individual sample, grouped by genotype. Colors represent gene-wise Z-score-transformed rlog expression values with red indicating high expression and blue indicating low expression. n = 9-10 G/G, 9-10 S/G, 8-9 S/S per tissue.

Taken together, RNA sequencing revealed multiple tissue-dependent differences in metabolic gene expression in PGC-1α^Ser481Gly^ mice, which may reflect cell-autonomous effects of the variants or adaptive response to systemic metabolic changes. While these transcriptional data do not establish causality, they suggest that coordinated alterations in oxidative gene expression across muscle, adipose tissue and liver may contribute to differences in fuel utilization and the response to metabolic challenges.

### Female S/S mice are more susceptible to weight gain and fat mass accumulation on chronic HFHS diet

To test whether mice carrying the serine allele could be more susceptible to energy storage under conditions of chronic dietary stress, we challenged male and female PGC-1α^Ser481Gly^ mice with a 14-week HFHS diet with *ad libitum* sugar-rich water, a model that promotes obesity and metabolic syndrome (57, 58). S/S females gained more weight compared to S/G and G/G mice which coincided with increased fat mass (Fig. 6a-c), consistent with the positive energy balance observed previously on this diet (Fig. 3m,p). In males, body weight and body composition were similar between genotypes, although S/S mice showed a trend toward reduced body weight gain over time compared to G/G mice (Fig. 6a-c). Together, these data further support that female S/S mice are more susceptible to body weight gain and fat mass accumulation under chronic dietary stress.

**Figure 6.**
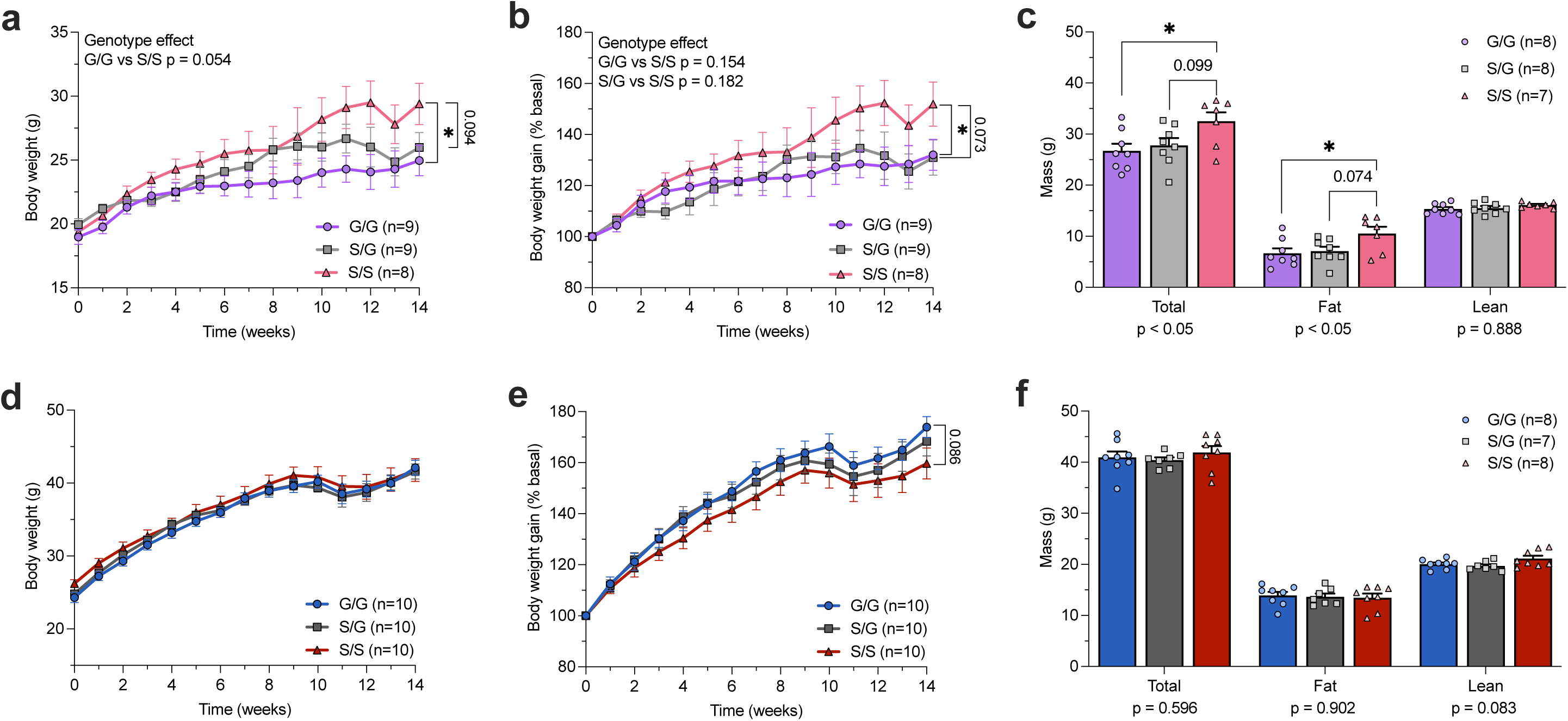
On a chronic HFHS diet, female S/S mice gain more weight and accumulate greater fat mass. **a,d**) Weekly body weight (g) and **b,e**) body weight gain (%) in female and male PGC-1α^Ser481Gly^ mice throughout the 14 weeks of HFHS diet combined with 30% fructose-rich water, respectively. **c,f**) Total body mass (g), fat mass (g), and lean mass (g) in female or male mice, respectively. Statistics: **a,b,d,e**) Two-way repeated measures ANOVA; **c,f**) Genotypes effects from one-way ANOVA, performed independently per body composition parameter. *p ≤ 0.05.

## Discussion

In this study, we investigated the effect of the PGC-1α Gly482Ser SNP on energy metabolism using cellular, physiological and transcriptomic approaches. In human iPSCs, the serine allele was associated with reduced cellular respiration, which was in line with decreased expression of genes in mitochondrial and fatty acid oxidative pathways in SCAT and liver of S/S mice. Intriguingly, these pathways were upregulated in S/S muscle, revealing an overall tissue-specific transcriptional reorganization of oxidative metabolism. Consistent with an overall shift toward anaerobic metabolism, indirect calorimetry revealed that S/S individuals living with overweight or obesity had increased postprandial carbohydrate oxidation following a high-fat meal. We observed a similar trend in female PGC-1α S/S mice, particularly in response to fasting. Female S/S mice were also prone to energy conservation during HFHS feeding, which manifested as reduced energy expenditure, positive energy balance and greater fat mass gain. Collectively, these results point toward a physiological and transcriptional predisposition toward impaired lipid oxidation and energy conservation in S/S females, leading to increases in body weight and fat mass following prolonged exposure to high caloric diets.

In this study, we observed increased carbohydrate oxidation rates, consistent with reduced lipid oxidation, in human carriers of the PGC-1α Gly482Ser SNP; however, a previous study in young, healthy adults found no differences in resting or maximal lipid oxidation between genotypes (59). Our data in mice suggest that impacts of the SNP are revealed in response to a metabolic challenge; thus, the above discrepancy may be explained by the absence of increased metabolic disease risk in their cohort. The stronger influence of the SNP in female mice also highlights sex as an additional modifier of PGC-1α-dependent metabolic regulation. Our data extend previous observations that the physiological consequences of altered PGC-1α activity are often sexually dimorphic, with female mice exhibiting greater susceptibility to hepatic oxidative damage, impaired brown adipose tissue thermogenesis and cardiovascular abnormalities in genetic models of reduced PGC-1α expression or activity (33, 55, 56). However, the absence of a dysfunctional metabolic phenotype in male mice carrying the S481 variant does not necessarily reflect the absence of metabolic risk in S/S males, particularly given the small sample size. Clinical studies have independently associated the serine allele with increased risk of T2D in males, increased risk of obesity in physically inactive males, and greater risk of hypertension (11, 13, 60). Of note, two of these studies reported their associations in males with a mean age above 50 years carrying a serine allele. Mice in the present study were approximately 22 weeks of age at the end of the 14-week diet period, corresponding to young adulthood (approximately 25-35 years of age) in humans. Together, these observations suggest that the metabolic consequences of this genetic variant are highly context-dependent and may emerge only in the presence of specific biological and environmental modifiers, such as sex, aging, obesity and physical inactivity.

Along these lines, fasting and HFHS feeding likely maximized our ability to detect biological and physiological differences in PGC-1α^Ser481Gly^ mice. Fasting is particularly informative in this context, as it requires a rapid transition from carbohydrate to lipid utilization, suggesting that the biological consequences of the serine allele are most apparent during conditions that demand PGC-1α-mediated metabolic adaptation. Consistent with this, glucagon, a key hormonal mediator of the fasting response, induced a smaller increase in PGC-1α expression in hepatocytes from female mice carrying a serine allele. Together with the reduced PGC-1α protein half-life reported for the S482 variant (33), these findings suggest that females carrying a serine allele may have a reduced capacity to upregulate PGC-1α in response to fasting, which could blunt adaptive metabolic responses, limiting lipid oxidation during macronutrient deprivation and favoring energy conservation during periods of nutrient excess. Supporting this, on a HFHS diet, S/S female mice showed lower energy expenditure, positive energy balance and greater fat mass, consistent with reports of lower metabolic rate and increased adiposity in females carrying PGC-1α variants (21, 61), and in female PGC-1α null mice (50). This propensity toward energy conservation is also evident in human isogenic S/S knock-in adipocytes, which exhibit increased lipogenesis and triacylglycerol accumulation (38). In line with a reduced PGC-1α activity model, adipocyte-specific overexpression of PGC-1α has the opposite effects, reducing adiposity and lipid droplet size while enhancing mitochondrial function and limiting hepatic lipid accumulation (62). These findings support a role for PGC-1α in regulating whole-body lipid partitioning and storage.

Because PGC-1α coordinates energy metabolism across multiple metabolically active tissues (15, 63), we supposed that the effects of this SNP would extend beyond adipose tissue and involve coordinated metabolic adaptations across organs. Consistent with this concept, mitochondrial and fatty acid oxidative pathways were downregulated in both SCAT and liver, suggesting a reduced capacity for lipid utilization and turnover in tissues central to the control of energy and lipid homeostasis. In contrast, these pathways were upregulated in muscle, which accounts for the majority of fatty acid oxidation during fasting (64, 65). The lack of a concomitant increase in whole-body lipid oxidation in S/S mice suggests that these transcriptional responses may be compensatory in nature. In iPSCs, we found that the serine allele translated into overall reduced oxidative activity; thus, the transcriptional induction of oxidative pathways in muscle may represent an adaptive response attempting to compensate for a cell-autonomous reduction in oxidative capacity. Several human studies suggest that PGC-1α serine allele carriers have impaired skeletal muscle oxidative function. This allele is associated with lower aerobic fitness, reduced VO_2_ max, and blunted muscle fibers adaptation to exercise training (66–68). Muscle PGC-1α expression is also reduced in elderly S/S carriers, individuals living with T2D, and those with impaired glucose tolerance and a family history of T2D (36, 12, 69), suggesting that PGC-1α S482 allele may predispose individuals toward reduced skeletal muscle oxidative capacity in response to metabolic challenges.

We have previously shown that loss of PGC-1α expression in skeletal muscle promotes systemic inflammation in liver and adipose tissue in elderly mice (70). These observations are consistent with the enrichment of inflammatory pathways in SCAT and liver of S/S mice, as well as reports linking the serine allele with heightened inflammatory responses (71). It was interesting that the most significantly affected gene linked to the SNP in our model was adipose *Tirap*, a protein that helps direct innate immunity defenses against pathogens (52). We did not pursue the potential implications of increased *Tirap* in the metabolic phenotypes, but it may reflect a novel link between PGC-1α and inflammation. Together, these findings suggest that the metabolic consequences of the serine allele may arise from coordinated alterations in oxidative metabolism and inflammatory signaling across muscle, SCAT and liver, two processes that are recognized hallmarks of metabolic disease (72–76). Direct functional assessment of oxidative capacity, lipid storage and inflammatory status in metabolic tissues are required to determine whether the transcriptional reprogramming observed in PGC-1α^Ser481Gly^ mice translates into functional impairments that could increase susceptibility to metabolic disease.

Beyond mechanistic characterization, our findings have potential implications for genotype-informed nutritional and exercise interventions aimed at preventing energy conservation and metabolic disease in S/S carriers. Since intermittent fasting relies precisely on a fasting-induced shift toward lipid oxidation to exert its metabolic benefits (77), it is interesting to speculate that S/S individuals may not derive the same metabolic benefits from this dietary approach. Physical activity may also represent a particularly relevant strategy to combat metabolic disease in serine allele carriers, since exercise is a significant inducer of PGC-1α expression and has been shown to prevent or reverse the metabolic abnormalities of reduced PGC-1α activity (78–82). Consistent with this, female T allele carriers of another PGC-1α SNP (rs17650401 C/T) who were following a hypocaloric diet had more significant fat loss following a 3-month fitness program (83). Exercise training also prevents weight gain induced by a high-fat high-fructose feeding in liver-specific PGC-1α KO mice (84). From a translational perspective, the PGC-1α Gly482Ser SNP could represent a valuable genetic marker to identify individuals, particularly females, who are at greater risk of metabolic dysfunction and may benefit from personalized lifestyle interventions.

Taken together, our data suggest that the serine allele reprograms whole-body metabolism toward an energy conversation phenotype, especially under certain dietary conditions. If the serine allele promotes energy conservation, its maintenance in human populations may be consistent with Neel’s thrifty genotype hypothesis (85), whereby genetic variants that favor efficient fuel storage confer a survival advantage during periods of food scarcity. In this context, the serine allele may be advantageous during periods of famine and cold exposure. Such selective pressures are particularly relevant in East Asian populations, which historically experienced repeated cycles of famine and exhibit the highest global frequency of the serine allele (19, 86). However, in the context of modern food abundance and physical inactivity, this once-beneficial genetic variant may now increase susceptibility to metabolic disease.

## Conclusion

In conclusion, our findings identify a novel role for the PGC-1α Gly482Ser SNP in regulating postprandial macronutrient oxidation and tissue-specific reprogramming of mitochondrial and fatty acid beta-oxidation metabolism across muscle, subcutaneous adipose tissue and liver in both sexes. Importantly, our data suggest that S/S female mice are predisposed to higher risk of energy conservation, linked to alterations in mitochondrial and oxidative gene programs across tissues and reduced energy expenditure. Beyond characterizing the metabolic consequences of this SNP, our findings in humans and mice reveal a broader role for PGC-1α in regulating metabolic adaptation to nutritional transitions.

## Limitations

A recognized challenge associated with our study of this SNP in humans was that low power affected our ability to detect statistically significant differences between genotypes. This was confounded by the lower frequency of the S/S genotype (10-15%) combined with the post-hoc nature of the analysis. Since the higher carbohydrate oxidation phenotype was not reproduced in S/S male mice, whether S/S males indeed have increased carbohydrate oxidation rates requires confirmation in a larger cohort. The transcriptomic analyses were limited by sample size per tissue, precluding the use of FDR-adjusted p-values for analysis. Therefore, we applied an unadjusted p-value threshold (p ≤ 0.05) for gene selection and pooled male and female PGC-1α^Ser481Gly^ mice to maximise statistical power. In this case, known tissue-specific sex difference in gene expression undoubtedly affected our ability to identify specific genes or pathways, increasing our chances of false negatives.

Although many interpretations were drawn cautiously from trends, the consistency of our findings across species strengthens their biological relevance and supports the existence of a genuine (albeit modest) genotype-dependent metabolic phenotype. This modest effect is consistent with the SNP acting as a disease-modifying rather than a disease-driving SNP. Its impact on metabolic disease risk may be potentiated by interactions with additional genetic variants, such as the PPARG Pro12Ala and PNPLA3 Ile148Met SNPs (87, 88), or by other environmental factors such as physical inactivity and diet.

## Methods

### Sex as a biological variable

Sex was included as a biological variable whenever feasible. Isogenic iPSC lines were derived from a single heterozygous male donor. In the human cohort, male and female participants were pooled due to insufficient statistical power for sex-disaggregated analyses. In mouse experiments, both sexes were included and data were analyzed separately, except for transcriptomic analyses, where male and female mice were pooled across genotypes to maximize statistical power.

### iPSCs cell lines and genome editing

Heterozygous (PGC1A c.1444G>A) 1-21 human iPSCs (89) were genome edited as previously described by the Tissue and Disease Modelling Core at BC Children’s Hospital Research Institute (90). Briefly the gRNA sequence (AGACAAGACCGGTGAACTGA) was cloned into pCCC as previously described (91). Five million iPSCs, grown in StemFlex (ThermoFisher, A3349401) were electroporated (Biorad GenePulser; 250V, 500µF) with 15 µg of pCCC and 15 µl of 125nt ssODN (20 µM) carrying either the G or A allele, a mutated PAM sequence, and a SacI site for genotyping. Cells were plated onto a Geltex-coated (ThermoFisher, A1413302) 60 mm plate (Falcon) and grown for 48 hours. Cells were then removed from plate with Accutase (StemCell Technologies), sorted for GFP (carried by pCCC), and then plated at clonal dilutions on Laminin521 (BioLamina) in StemFlex. After 7 days, colonies were picked onto Laminin521-coated 96 well plates, grown for 4 days, replica plated and then frozen for storage or used for PCR genotyping (AGGAAAGGAAGACCAAGCGG; AAGGACCTTGAACGAGAGCG) and SacI digest. Positive clones were confirmed by sequencing and then expanded into 12-well plates prior to re-genotyping and freezing.

### Study participants and clinical

The clinical data analyzed in this study were obtained from research volunteers who were free of chronic disease or medication affecting metabolism, who participated in two clinical trials between 2010-2020 by the team of May Faraj at the IRCM, as reported previously (47, 48). Participants who accepted to donate a DNA sample for research on genes related to lipid metabolism were included in this analysis. Participants were stratified by genotype into: G/G (N = 12; 4 males), S/G (N = 21; 7 males), and S/S (N = 5; 2 males). Detailed information regarding the study populations, clinical testing and sample analyses can be found in the original publications. Postprandial carbohydrate oxidation, respiratory quotient and energy expenditure were measured after an overnight 12-hour fast, following consumption of a high-fat meal (600 kcal/m^2^, 68% fat, 18% carbohydrate). Body composition was assessed by dual-energy Xray absorptiometry (GE Healthcare), basal metabolic rate and substrate oxidation by indirect calorimetry (Vmax Encore; Carefusion).

### Animals

Age-and sex-matched PGC-1α^Ser481Gly^ mice were housed under controlled conditions (22°C, 12-hour light/dark cycle) with *ad libitum* access to food and water. Starting at 8 weeks of age, mice were assigned either to a chow diet (D12450KI; 10% kcal from fat, Research Diets Inc.) with regular water or, in a separate cohort, to a high-fat high-sugar (HFHS) diet (D12451I; 60% kcal from fat, Research Diets Inc.) combined with 30% D-fructose in drinking water. Each cohort of mice was maintained on its respective diet for 14 weeks.

### Generation of mice harboring the 481Gly allele

Two guide RNAs (gRNA) specific for the desired *PPARGC1A* 1441 A>G mutation, converting the serine-encoding A to the glycine-encoding G nucleotides on the coding strand, were cloned into the Cas9-encoding px330 plasmid (Guide 1: 5’ – CAGACAAGACCAGTGAACTA – 3’ and Guide 2: 5’ – AAGACCAGTGAACTAAGGGA – 3’). Embryos from B6C3F1 zygotes were microinjected (IRCM core facilities) with the two plasmids together, each expressing a single guide with the single-stranded oligodeoxynucleotides (ssODNs) (5’ – TGAACAAGCACTTCGGTCATCCCTGTCAAGCTGTGTTTGACGACAAATCAGACAAG ACCGGTGAACTAAGAGATGGCGACTTCAGTAATGAACAATTCTCCAAACTACCTGTGTTTA TAA – 3’). The embryos were implanted into pseudo-pregnant female mice. After weaning, 500 pups (F0) were screened using a Taqman SNP genotyping kit and the presence of the PGC-1α 481Gly and 481Ser alleles was verified using Sanger Sequencing. Two males and two females were backcrossed with C57BL/6N wildtype mice to ensure transmission of the allele would not be affected by sex. The resulting offspring (F1) were screened using PCR (F: 5’-GCTAATGGATCCTACATTTCTTTTTGTTTC -3’ and R: 5’-GCAACTTGCCTCTTAGCGC-3’) and subsequent Age1 digestion. Further genotyping was performed using a KASP genotyping assay kit (KBS-1050-121, LGC Genomics KBiosciences) according to the manufacturer’s standard protocol. Genotype frequencies did not deviate from the expected Mendelian 1:2:1 ratio.

### Body weight imputation and cohort normalization

For longitudinal body weight analyses on a chow diet, two cohorts of mice were pooled. Cohort-to-cohort technical variation and missing weekly body weight measurements were addressed through a two-step procedure. First, missing body weight values were imputed using a linear mixed-effects model with time, genotype, and their interaction as fixed effects, and mouse-specific random intercepts and slopes to account for individual growth trajectories. Second, cohort effects were removed by fitting, at each week, a linear model with cohort as the predictor; residuals were then added back to the overall weekly mean to generate cohort-adjusted body weights. This normalization was applied to both observed and imputed values. Mice that died before the study endpoint were excluded prior to any body weight imputation or normalization.

### Indirect calorimetric measurements

Metabolic measurements were performed using the Promethion metabolic cages (Sable Systems International) after 14 weeks of diet. Male and female mice aged 22 weeks were individually housed in metabolic cages at 22 °C under a 12-h light–dark cycle. After the initial 72 hours of acclimation in the cage, the SableScreen software (v3.3.12.0) recorded cage and metabolic activity and condensed the data into 30-minute intervals. All time-series data are plotted as group mean ± SEM, smoothed using a 6-point centered rolling mean (3-h window) for visualization purposes only. Mice were maintained with *ad libitum* access to food and water during the entire measurement, except during the 24-h fast. Cage activity was measured as the cumulative AllMeters parameter generated from the Promethion X-Y-Z infrared beam-break array, which include ambulatory and non-ambulatory movements, such as grooming. Voluntary exercise was assessed by providing mice with free access to a running wheel within the metabolic cages. Metabolic activity was measured via indirect calorimetry recording maximal O_2_ consumption (VO_2_) and CO_2_ production (VCO_2_). RER was calculated as VCO_2_/VO_2_. Energy expenditure was calculated from the Weir equation (92). Food caloric intake (kcal) was calculated by multiplying food consumed (g) by the metabolizable energy content of the diet (3.82 kcal/g for chow and 4.73 kcal/g for HFHS). Water intake was recorded in grams and converted to milliliters using the density of water (1 g/mL). Energy balance was calculated as food intake minus energy expenditure. Average rates were calculated for each animal during the dark (18:00–06:00) and light (06:00–18:00) phases over the specified period. A combined-phase value was then calculated as the mean of the dark- and light-phase averages to ensure equal weighting of both phases. All analyses were performed in RStudio (version 2026.05). Following quality control, animals with missing data resulting from death or cage malfunction were excluded from the relevant analyses. Sample sizes for each analysis are available in the figures. For further information, please refer to the metadata tables (Supplementary File 4).

### Magnetic resonance imaging

Whole-body lean and fat mass were measured by magnetic resonance imaging (MRI) using the minispec mq 7.5 NMR analyzer (Bruker, MA) on animals fed 14-weeks of chow or HFHS diet. Live animals were weighed and their ear-tags were removed before MRI. Then, protocol was followed according to manufacturer’s instructions to obtain fat and lean mass data. For ANCOVA analysis, mean body weight measured immediately before and after the metabolic cage housing period was used as a covariate.

### RNA extraction, 3′ digital gene expression library preparation and sequencing

The same group of PGC-1α^Ser481Gly^ mice fed a chow diet (D12450KI; Research Diets Inc.) for 14 weeks were maintained on the same diet for a total of 24 weeks. Mice were fasted for 3 hours before tissue collection. RNA was isolated from ∼10 mg liver, ∼40 mg SCAT and ∼20 mg gastrocnemius muscle tissue (n=10-11 per genotype and per tissue) using TRIzol reagent (Invitrogen #15596018) and concentrated with RNA Clean & Concentrator-25 kit (Zymo Research #R1018, USA). The RNA concentration was quantified using NanoDrop Spectrophotometer (NanoDrop Technologies), and the integrity (RIN score >7) was assessed with the Agilent RNA 6000 Nano kit on the Agilent Bioanalyzer 2100.

Transcriptome profiling was performed using a 3′ digital gene expression sequencing (3′ DGE-seq) protocol based on the 3′ Sequencing RNA Profiling (3′SRP) approach (*93, 94*). Libraries were prepared from 10 ng of total RNA per sample. During template-switching reverse transcription, polyadenylated RNA molecules were tagged with universal adapters, sample-specific barcodes, and unique molecular identifiers (UMIs). Barcoded cDNAs from individual samples were pooled and amplified, followed by transposase-mediated tagmentation to generate libraries enriched for the 3′ ends of transcripts. Libraries were sequenced on an Illumina NovaSeq 6000 platform, yielding an average of approximately 5 million 75-bp transcript reads per sample.

### DGE-seq processing and differential expression analysis

Sequencing data were processed using the 3′ Sequencing RNA Profiling (SRP) pipeline. Briefly, reads were demultiplexed according to sample-specific barcodes, and transcript reads were aligned to the mouse RefSeq reference transcriptome corresponding to the mm10 genome assembly using BWA. Reads mapping ambiguously to genes or failing the pipeline quality criteria were excluded, and gene-level expression matrices were generated by counting unique UMIs assigned to each gene.

Sample-level quality control was based on the number of detected genes and total UMI counts. Genes with a summed UMI count ≤150 across all samples were excluded prior to downstream analyses. Filtered raw UMI counts were used as input for differential expression analysis, whereas regularized-logarithm (rlog)-transformed counts were used for principal component analysis (PCA) and data visualization. Outlier detection was performed separately for each tissue using Mahalanobis distances calculated from the first two principal components. Samples exceeding the 97.5th percentile of the corresponding chi-squared distribution were classified as outliers, resulting in the exclusion of seven samples from subsequent analyses.

Differential gene expression was assessed independently in muscle, SCAT and liver using DESeq2, with sex included as a covariate in the model (∼ sex + genotype + sex:genotype). Genes with a nominal P < 0.05 and an absolute log2 fold change >1 were considered differentially expressed. For downstream pathway analyses, genes were ranked according to their DESeq2 Wald test statistics. Rlog-transformed expression values were used for visualization.

### Gene set enrichment and pathway activity analyses (GSEA)

GSEA was performed using the fgsea R package and Gene Ontology Biological Process gene sets obtained from the Molecular Signatures Database (MSigDB, v2024.1, mouse). Genes were ranked by their DESeq2 Wald statistics, and enrichment was evaluated using 1,000 permutations. Pathways with a nominal P < 0.05 were retained for exploratory interpretation. Shared and tissue-specific transcriptional responses were identified by comparing enriched pathways across muscle, SCAT and liver. For selected pathways, leading-edge genes identified by fgsea were ranked according to their absolute DESeq2 Wald statistic, and the 15 highest-ranking genes were visualized using gene-wise Z-score-transformed rlog expression values.

Estrogen receptor signaling activity was assessed by ssGSEA using the GSVA R package and the GOBP_ESTROGEN_RECEPTOR_SIGNALING_PATHWAY gene set. ssGSEA scores were calculated separately from rlog-transformed expression data for each tissue. Differences in pathway activity between genotypes were assessed using limma. Analyses combining both sexes included sex as a covariate (∼ sex + genotype), whereas sex-stratified analyses included genotype alone (∼ genotype). Statistical inference was based on limma empirical Bayes-moderated statistics.

### Oxygen consumption rate

iPSCs grown in mTeSR™ Plus medium (STEMCELL Technologies, 100-0276) were dissociated using TrypLE™ (Thermofisher, 12604021) and seeded at a density of 7-11 x10^3^ cells per well onto 96-well flat-bottom plates, pre-coated with Matrigel (Corning, 354277), in a final volume of 200 μl of mTeSR medium containing 10 μM Y-27632 (STEMCELL Technologies, 72302). Cell culture media was replenished 24 hours later with 100 μl of mTeSR and oxygen consumption rates (OCR) were monitored in real time using the Resipher platform equipped with a 32-sensor lid (Lucid Scientific, #NS32-101A) at 37°C and 5% CO_2_. All unused wells were filled with an equivalent volume of PBS. When basal OCR reached a plateau, iPSCs were treated with 0.125 μM Oligomycin (Abcam, Ab141829,) or 4 μM FCCP (Abcam, Ab120081) in mTeSR Plus medium. After 4 hours, FCCP-treated wells were replenished with medium containing 1.5 μM rotenone (Abcam, Ab143145) and monitored for an additional 3 hours. DMSO-treated wells served as the vehicle controls. Each condition was assayed in 5-10 technical replicate wells per clone. OCR values were calculated using Lucid Lab software and normalized to the mean cellular protein content. Total cellular protein content was quantified from parallel wells harvested at the beginning of the experiment in 20 μl RIPA lysis buffer using DC Protein Assay Reagents (Biorad, 5000113-4). Basal respiration was calculated as the difference between OCR in DMSO-treated and rotenone-treated wells. ATP-linked respiration was defined as the difference between DMSO-treated and oligomycin-treated wells. Spare respiratory capacity was calculated as the difference between FCCP-treated and DMSO-treated wells, whereas maximal respiration was calculated as the difference between FCCP-treated and rotenone-treated wells. Proton leak was determined as the difference between oligomycin-treated and rotenone-treated wells. Non-mitochondrial respiration was defined as OCR measured in rotenone-treated wells following correction for background oxygen consumption in cell-free wells.

### Primary hepatocyte isolation and Western Blotting

Primary mouse hepatocytes were isolated according to established protocols (95) from 12-to 17-week-old male and female PGC-1α^Ser481Gly^ mice fed a standard laboratory chow (Teklad Global 18% Protein Rodent Diet 2018; Envigo). Cells were plated at a density of 0.45 x 10^6^ cells/well in 6-well plates. Following a 2-hour recovery period in plating medium consisting of DMEM supplemented with 4.5 g/L glucose (Multicell, 319-005-CL), 10% fetal bovine serum, 2 mM sodium pyruvate, 1% penicillin-streptomycin, 1 μM dexamethasone, and 100 nM insulin, cells were incubated for 16 hours in starvation medium consisting of DMEM supplemented with 1.0 g/L glucose (Multicell, 319-010-CL), 0.2% bovine serum albumin (BSA) Fraction V, 2 mM sodium pyruvate, and 1% penicillin-streptomycin. For glucagon treatment, cells were incubated in starvation medium supplemented with 50 nM glucagon (Cayman Chemical, 24204) for 3 hours. Total cell lysates were prepared using RIPA buffer and analyzed by western blot using antibodies against PGC-1α (Millipore, ST1202), beta-actin (Sigma, A5441) and Hsp90 (Cell Signaling, 4874S).

### Statistics and reproducibility

Outliers were identified using the ROUT method (Q=1%) and quality control plots. Normality was assessed using the Shapiro–Wilk test where applicable. An unpaired Welch’s t-test was used when comparing two groups. For repeated measures comparing two or more groups over time, a two-way repeated measures ANOVA with Geisser-Greenhouse correction was performed, followed by post hoc analyses as indicated in the figure legends. For independent comparisons of three groups with one variable, including mass-independent parameters (RER, energy balance and exercise), either a one-way ANOVA, Welch’s ANOVA or a Kruskal-Wallis test was employed. For mass-dependent parameters, ANCOVA was used unless a significant genotype × body mass interaction was detected; in which case a generalized linear model (GLM) including the interaction term was used. For all indirect calorimetry parameters, statistical analyses were performed between genotypes independently for each diet or photoperiod (combined, dark, light). Pairwise genotype comparisons were performed using Tukey-adjusted post hoc tests. Data are expressed as mean ± SEM. Statistical analyses were performed using Prism (version 10, GraphPad Software Inc) and R (version 2026.05). Statistical significance was defined as p ≤ 0.05. Results that did not reach statistical significance but were considered biologically relevant are described as trends.

### Study approval

This study was approved by the IRCM Research Ethics Board (REB) (research participants: # 2017-903; iPSC studies: 2020-1047), the BC Children’s Hospital REB (H09-00676), and the IRCM Animal Care Committee (2022–02). Written informed consent was obtained from all human participants.

### Data availability

The data supporting the findings of this study are documented within the paper and are available from the corresponding authors upon request. Correspondence and requests for materials should be addressed to: Jennifer L. Estall.

## Author Contributions

MG performed *in vitro* experiments, analyzed and interpreted data, and wrote the manuscript. NJ and EC designed the study and performed animal experiments. RV and JLE generated the PGC-1α^Ser481Gly^ mouse model. CF, MB and JP performed the RNA-seq analysis. MS prepared figures and contributed to the RNA-seq bioinformatic analysis. FCL generated the human iPSC lines. MF provided clinical data and analysis. JLE conceptualized and financially supported the study. AG and JLE contributed to manuscript writing and editing. All authors reviewed and approved the manuscript.

## AI disclosure

Microsoft Copilot (M365 Copilot, based on the GPT-5 chat model) was used solely for language and grammatical editing. All AI-generated revisions were carefully reviewed by the authors to ensure that the presentation and interpretation of the data were not impacted. AI tools were not used to generate figures, analyze data, or draw conclusions. The authors take full responsibility for the content of the manuscript.

## Acknowledgments

We extend our gratitude to the animal facility staff for their diligent care of the mice used in this study.

---

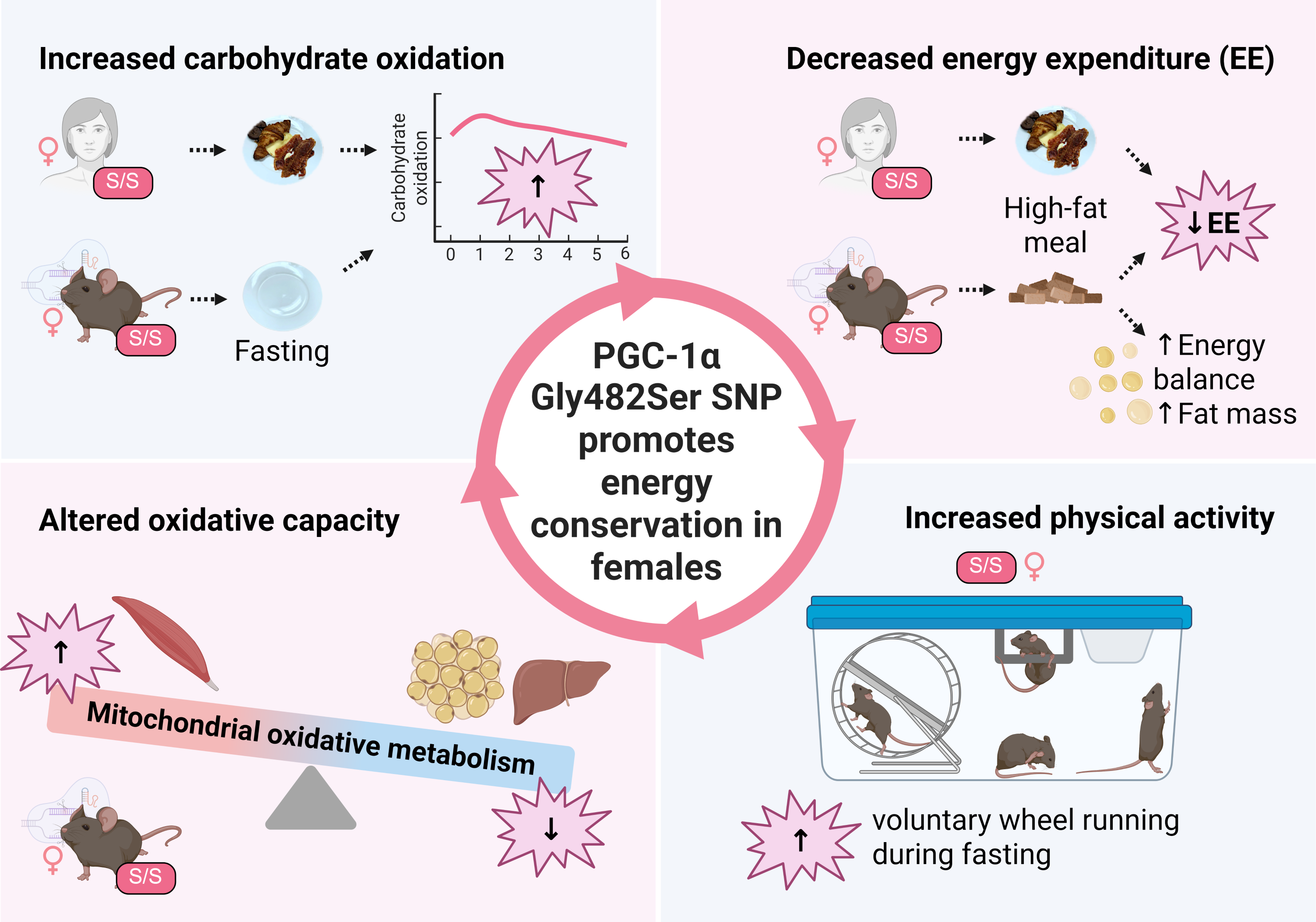

## References

1. Almgren P, et al. Heritability and familiality of type 2 diabetes and related quantitative traits in the Botnia Study. Diabetologia. 2011;54(11):2811–2819.

2. Elks CE, et al. Variability in the Heritability of Body Mass Index: A Systematic Review and Meta-Regression. Front Endocrinol. 2012;3. 10.3389/fendo.2012.00029.

3. Willemsen G, et al. The Concordance and Heritability of Type 2 Diabetes in 34,166 Twin Pairs From International Twin Registers: The Discordant Twin (DISCOTWIN) Consortium. Twin Research and Human Genetics. 2015;18(6):762–771.

4. Bouchard C, et al. The response to long-term overfeeding in identical twins. N Engl J Med. 1990;322(21):1477–1482.

5. Ravussin E, et al. Effects of a traditional lifestyle on obesity in Pima Indians. Diabetes Care. 1994;17(9):1067–1074.

6. Mahajan A, et al. Genome-wide trans-ancestry meta-analysis provides insight into the genetic architecture of type 2 diabetes susceptibility. Nat Genet. 2014;46(3):234–244.

7. Locke AE, et al. Genetic studies of body mass index yield new insights for obesity biology. Nature. 2015;518(7538):197–206.

8. Wild S, et al. Global Prevalence of Diabetes: Estimates for the year 2000 and projections for 2030. Diabetes Care. 2004;27(5):1047–1053.

9. Abarca-Gómez L, et al. Worldwide trends in body-mass index, underweight, overweight, and obesity from 1975 to 2016: a pooled analysis of 2416 population-based measurement studies in 128·9 million children, adolescents, and adults. The Lancet. 2017;390(10113):2627–2642.

10. Oberkofler H, et al. Peroxisome proliferator-activated receptor-gamma coactivator-1 gene locus: associations with hypertension in middle-aged men. Hypertension. 2003;41(2):368– 372.

11. Cheurfa N, et al. The Gly482Ser polymorphism in the peroxisome proliferator-activated receptor-gamma coactivator-1 gene is associated with hypertension in type 2 diabetic men. Diabetologia. 2004;47(11):1980–1983.

12. Ling C, et al. Multiple environmental and genetic factors influence skeletal muscle PGC-1α and PGC-1β gene expression in twins [Internet]. 2004. 10.1172/JCI21889.

13. Ridderstråle M, et al. Increased risk of obesity associated with the variant allele of the PPARGC1A Gly482Ser polymorphism in physically inactive elderly men. Diabetologia. 2006;49(3):496–500.

14. Okauchi Y, et al. PGC-1alpha Gly482Ser polymorphism is associated with the plasma adiponectin level in type 2 diabetic men. Endocr J. 2008;55(6):991–997.

15. Puigserver P, et al. A cold-inducible coactivator of nuclear receptors linked to adaptive thermogenesis. Cell. 1998;92(6):829–839.

16. Wu Z, et al. Mechanisms Controlling Mitochondrial Biogenesis and Respiration through the Thermogenic Coactivator PGC-1. Cell. 1999;98(1):115–124.

17. Lehman JJ, et al. Peroxisome proliferator-activated receptor gamma coactivator-1 promotes cardiac mitochondrial biogenesis. J Clin Invest. 2000;106(7):847–856.

18. Vega RB, Huss JM, Kelly DP. The Coactivator PGC-1 Cooperates with Peroxisome Proliferator-Activated Receptor α in Transcriptional Control of Nuclear Genes Encoding Mitochondrial Fatty Acid Oxidation Enzymes. Molecular and Cellular Biology. 2000;20(5):1868–1876.

19. rs8192678 RefSNP Report - dbSNP - NCBI [Internet]. 2026. https://www.ncbi.nlm.nih.gov/snp/rs8192678#frequency_tab. Accessed July 21, 2026.

20. Ek J, et al. Mutation analysis of peroxisome proliferator-activated receptor-γ coactivator-1 (PGC-1) and relationships of identified amino acid polymorphisms to Type II diabetes mellitus. Diabetologia. 2001;44(12):2220–2226.

21. Esterbauer H, et al. Peroxisome Proliferator-Activated Receptor-γ Coactivator-1 Gene Locus: Associations With Obesity Indices in Middle-Aged Women. Diabetes. 2002;51(4):1281–1286.

22. Hara K, et al. A genetic variation in the PGC-1 gene could confer insulin resistance and susceptibility to Type II diabetes. Diabetologia. 2002;45(5):740–743.

23. Deeb SS, Brunzell JD. The Role of the PGC1α Gly482Ser Polymorphism in Weight Gain due to Intensive Diabetes Therapy. PPAR Res. 2009;2009:649286.

24. Xia W, et al. Systematic Meta-analysis Revealed an Association of PGC-1α rs8192678 Polymorphism in Type 2 Diabetes Mellitus. Dis Markers. 2019;2019:2970401.

25. Ling C, et al. Epigenetic regulation of PPARGC1A in human type 2 diabetic islets and effect on insulin secretion. Diabetologia. 2008;51(4):615–622.

26. Zhang R-N, et al. PPARGC1A rs8192678 G>A polymorphism affects the severity of hepatic histological features and nonalcoholic steatohepatitis in patients with nonalcoholic fatty liver disease. World J Gastroenterol. 2021;27(25):3863–3876.

27. Yoon JC, et al. Control of hepatic gluconeogenesis through the transcriptional coactivator PGC-1. Nature. 2001;413(6852):131–138.

28. Baar K, et al. Adaptations of skeletal muscle to exercise: rapid increase in the transcriptional coactivator PGC-1. The FASEB Journal. 2002;16(14):1879–1886.

29. Suliman HB, et al. Lipopolysaccharide induces oxidative cardiac mitochondrial damage and biogenesis. Cardiovasc Res. 2004;64(2):279–288.

30. Handschin C, et al. Nutritional Regulation of Hepatic Heme Biosynthesis and Porphyria through PGC-1α. Cell. 2005;122(4):505–515.

31. Sano M, et al. Intramolecular Control of Protein Stability, Subnuclear Compartmentalization, and Coactivator Function of Peroxisome Proliferator-activated Receptor γ Coactivator 1α *. Journal of Biological Chemistry. 2007;282(35):25970–25980.

32. Palomer X, et al. TNF-α reduces PGC-1α expression through NF-κB and p38 MAPK leading to increased glucose oxidation in a human cardiac cell model. Cardiovasc Res. 2009;81(4):703–712.

33. Besse-Patin A, et al. Estrogen Signals Through Peroxisome Proliferator-Activated Receptor-γ Coactivator 1α to Reduce Oxidative Damage Associated With Diet-Induced Fatty Liver Disease. Gastroenterology. 2017;152(1):243–256.

34. Chen Y, et al. Gly482Ser mutation impairs the effects of peroxisome proliferator–activated receptor *γ* coactivator-1*α* on decreasing fat deposition and stimulating phosphoenolpyruvate carboxykinase expression in hepatocytes. Nutrition Research. 2013;33(4):332–339.

35. Taghvaei S, Saremi L, Babaniamansour S. Computational Analysis of Gly482Ser Single-Nucleotide Polymorphism in PPARGC1A Gene Associated with CAD, NAFLD, T2DM, Obesity, Hypertension, and Metabolic Diseases. PPAR Res. 2021;2021:5544233.

36. Patti ME, et al. Coordinated reduction of genes of oxidative metabolism in humans with insulin resistance and diabetes: Potential role of PGC1 and NRF1. Proceedings of the National Academy of Sciences. 2003;100(14):8466–8471.

37. Semple RK, et al. Expression of the thermogenic nuclear hormone receptor coactivator PGC-1alpha is reduced in the adipose tissue of morbidly obese subjects. Int J Obes Relat Metab Disord. 2004;28(1):176–179.

38. Huang M, et al. Engineered allele substitution at PPARGC1A rs8192678 alters human white adipocyte differentiation, lipogenesis, and PGC-1α content and turnover. Diabetologia. 2023;66(7):1289–1305.

39. Muller YL, et al. A Gly482Ser Missense Mutation in the Peroxisome Proliferator-Activated Receptor γ Coactivator-1 Is Associated With Altered Lipid Oxidation and Early Insulin Secretion in Pima Indians. Diabetes. 2003;52(3):895–898.

40. Stefan N, et al. Genetic Variations in PPARD and PPARGC1A Determine Mitochondrial Function and Change in Aerobic Physical Fitness and Insulin Sensitivity during Lifestyle Intervention. J Clin Endocrinol Metab. 2007;92(5):1827–1833.

41. Ramos-Lopez O, et al. Association of the Gly482Ser PPARGC1A gene variant with different cholesterol outcomes in response to two energy-restricted diets in subjects with excessive weight. Nutrition. 2018;47:83–89.

42. Zhang Y, et al. The rs17782313 polymorphism near MC4R gene confers a high risk of obesity and hyperglycemia, while PGC1α rs8192678 polymorphism is weakly correlated with glucometabolic disorder: a systematic review and meta-analysis. Front Endocrinol. 2023;14. 10.3389/fendo.2023.1210455.

43. Smith RL, et al. Metabolic Flexibility as an Adaptation to Energy Resources and Requirements in Health and Disease. Endocr Rev. 2018;39(4):489–517.

44. Palmer BF, Clegg DJ. Metabolic Flexibility and Its Impact on Health Outcomes. Mayo Clin Proc. 2022;97(4):761–776.

45. Bakkman L, et al. Reduced Respiratory Capacity in Muscle Mitochondria of Obese Subjects. Obes Facts. 2010;3(6):371–375.

46. Chen T-H, et al. Mitochondrial Dysfunction as an Underlying Cause of Skeletal Muscle Disorders. Int J Mol Sci. 2022;23(21):12926.

47. Bissonnette S, et al. High plasma apolipoprotein B identifies obese subjects who best ameliorate white adipose tissue dysfunction and glucose-induced hyperinsulinemia after a hypocaloric diet. Am J Clin Nutr. 2018;108(1):62–76.

48. Bissonnette S, et al. Native low-density lipoproteins are priming signals of the NLRP3 inflammasome/interleukin-1β pathway in human adipose tissue and macrophages. Sci Rep. 2023;13(1):18848.

49. Lin J, et al. Defects in Adaptive Energy Metabolism with CNS-Linked Hyperactivity in PGC-1α Null Mice. Cell. 2004;119(1):121–135.

50. Leone TC, et al. PGC-1alpha deficiency causes multi-system energy metabolic derangements: muscle dysfunction, abnormal weight control and hepatic steatosis. PLoS Biol. 2005;3(4):e101.

51. Elia M. Organ and tissue contribution to metabolic rate. In: Kinney JM, Tucker HN, eds. Energy Metabolism: Tissue Determinants and Cellular Corollaries. New York, NY: Raven Press; 1992:61–80.

52. Horng T, Barton GM, Medzhitov R. TIRAP: an adapter molecule in the Toll signaling pathway. Nat Immunol. 2001;2(9):835–841.

53. Vandenbeek R, Khan NP, Estall JL. Linking Metabolic Disease With the PGC-1α Gly482Ser Polymorphism. Endocrinology. 2018;159(2):853–865.

54. Morselli E, et al. Hypothalamic PGC-1α Protects Against High-Fat Diet Exposure by Regulating ERα. Cell Reports. 2014;9(2):633–645.

55. Naumenko N, et al. PGC-1α deficiency reveals sex-specific links between cardiac energy metabolism and EC-coupling during development of heart failure in mice. Cardiovasc Res. 2022;118(6):1520–1534.

56. Takeuchi A, et al. Sex difference in BAT thermogenesis depends on PGC-1α–mediated phospholipid synthesis in mice. Nat Commun. 2025;16(1):6072.

57. Bray GA, Nielsen SJ, Popkin BM. Consumption of high-fructose corn syrup in beverages may play a role in the epidemic of obesity. Am J Clin Nutr. 2004;79(4):537–543.

58. Messier C, et al. The effects of a high-fat, high-fructose, and combination diet on learning, weight, and glucose regulation in C57BL/6 mice. Behavioural Brain Research. 2007;178(1):139–145.

59. Montes-de-Oca-García A, et al. Influence of Peroxisome Proliferator-Activated Receptor (PPAR)-gamma Coactivator (PGC)-1 alpha gene rs8192678 polymorphism by gender on different health-related parameters in healthy young adults. Front Physiol. 2022;13. 10.3389/fphys.2022.885185.

60. Sun L, et al. The Gly482Ser variant of the PPARGC1 gene is associated with Type 2 diabetes mellitus in northern Chinese, especially men. Diabetic Medicine. 2006;23(10):1085– 1092.

61. Moradi S, et al. Variants in the PPARGC1A Gene may Influence the Effect of Fat Intake on Resting Metabolic Rate in Obese Women. Lipids. 2018;53(3):291–300.

62. Shen S-H, et al. Adipocyte-Specific Expression of PGC1α Promotes Adipocyte Browning and Alleviates Obesity-Induced Metabolic Dysfunction in an HO-1-Dependent Fashion. Antioxidants (Basel). 2022;11(6):1147.

63. Finck BN, Kelly DP. PGC-1 coactivators: inducible regulators of energy metabolism in health and disease. J Clin Invest. 2006;116(3):615–622.

64. van Loon LJC, et al. The effects of increasing exercise intensity on muscle fuel utilisation in humans. The Journal of Physiology. 2001;536(1):295–304.

65. van Loon LJC, et al. Intramyocellular lipids form an important substrate source during moderate intensity exercise in endurance-trained males in a fasted state. The Journal of Physiology. 2003;553(2):611–625.

66. Akhmetov II, et al. [Association of regulatory genes polymorphisms with aerobic and anaerobic performance of athletes]. Ross Fiziol Zh Im I M Sechenova. 2007;93(8):837–843.

67. Steinbacher P, et al. The Single Nucleotide Polymorphism Gly482Ser in the PGC-1α Gene Impairs Exercise-Induced Slow-Twitch Muscle Fibre Transformation in Humans. PLOS ONE. 2015;10(4):e0123881.

68. Chen Y, et al. Meta-analyses of the association between the PPARGC1A Gly482Ser polymorphism and athletic performance. Biol Sport. 2019;36(4):301–309.

69. Barrès R, et al. Non-CpG Methylation of the PGC-1α Promoter through DNMT3B Controls Mitochondrial Density. Cell Metabolism. 2009;10(3):189–198.

70. Sczelecki S, et al. Loss of Pgc-1α expression in aging mouse muscle potentiates glucose intolerance and systemic inflammation. American Journal of Physiology-Endocrinology and Metabolism. 2014;306(2):E157–E167.

71. Pasquinelli A, et al. Gly482Ser PGC-1α Gene Polymorphism and Exercise-Related Oxidative Stress in Amyotrophic Lateral Sclerosis Patients. Front Cell Neurosci. 2016;10. 10.3389/fncel.2016.00102.

72. Pérez-Carreras M, et al. Defective hepatic mitochondrial respiratory chain in patients with nonalcoholic steatohepatitis. Hepatology. 2003;38(4):999–1007.

73. Dahlman I, et al. Downregulation of electron transport chain genes in visceral adipose tissue in type 2 diabetes independent of obesity and possibly involving tumor necrosis factor-alpha. Diabetes. 2006;55(6):1792–1799.

74. Ciupińska-Kajor M, et al. Hepatic angiogenesis and fibrosis are common features in morbidly obese patients. Hepatol Int. 2011;7(1):233–240.

75. Heinonen S, et al. Impaired Mitochondrial Biogenesis in Adipose Tissue in Acquired Obesity. Diabetes. 2015;64(9):3135–3145.

76. Bou Matar D, et al. Adipose tissue dysfunction disrupts metabolic homeostasis: mechanisms linking fat dysregulation to disease. Front Endocrinol (Lausanne). 2025;16:1592683.

77. Anton SD, et al. Flipping the Metabolic Switch: Understanding and Applying the Health Benefits of Fasting. Obesity. 2018;26(2):254–268.

78. Lanza IR, Nair KS. Muscle mitochondrial changes with aging and exercise. Am J Clin Nutr. 2009;89(1):467S–71S.

79. Hernández-Alvarez MI, et al. Subjects with early-onset type 2 diabetes show defective activation of the skeletal muscle PGC-1{alpha}/Mitofusin-2 regulatory pathway in response to physical activity. Diabetes Care. 2010;33(3):645–651.

80. Barrès R, et al. Acute exercise remodels promoter methylation in human skeletal muscle. Cell Metab. 2012;15(3):405–411.

81. Summermatter S, Handschin C. PGC-1α and exercise in the control of body weight. Int J Obes (Lond). 2012;36(11):1428–1435.

82. Ringholm S, et al. Effect of lifelong resveratrol supplementation and exercise training on skeletal muscle oxidative capacity in aging mice; impact of PGC-1α. Exp Gerontol. 2013;48(11):1311–1318.

83. Mazur II, et al. PPARGC1A gene polymorphism is associated with exercise-induced fat loss. Mol Biol Rep. 2020;47(10):7451–7457.

84. Kristensen CM, et al. PGC-1α in hepatic UPR during high-fat high-fructose diet and exercise training in mice. Physiological Reports. 2018;6(15):e13819.

85. Neel JV. Diabetes mellitus: a “thrifty” genotype rendered detrimental by “progress”? Am J Hum Genet. 1962;14(4):353–362.

86. Li LM. Introduction: Food, Famine, and the Chinese State. J of Asian Stud. 1982;41(4):687–707.

87. Ruchat S-M, et al. Evidence for interaction between PPARG Pro12Ala and PPARGC1A Gly482Ser polymorphisms in determining type 2 diabetes intermediate phenotypes in overweight subjects. Exp Clin Endocrinol Diabetes. 2009;117(9):455–459.

88. Tai C-M, et al. Interactions of a PPARGC1A Variant and a PNPLA3 Variant Affect Nonalcoholic Steatohepatitis in Severely Obese Taiwanese Patients. Medicine (Baltimore). 2016;95(12):e3120.

89. Krentz NAJ, et al. Phosphorylation of NEUROG3 Links Endocrine Differentiation to the Cell Cycle in Pancreatic Progenitors. Dev Cell. 2017;41(2):129–142.e6.

90. Frew J, et al. Premature termination codon readthrough upregulates progranulin expression and improves lysosomal function in preclinical models of GRN deficiency. Mol Neurodegener. 2020;15(1):21.

91. Krentz NAJ, Nian C, Lynn FC. TALEN/CRISPR-mediated eGFP knock-in add-on at the OCT4 locus does not impact differentiation of human embryonic stem cells towards endoderm. PLoS One. 2014;9(12):e114275.

92. Weir JB de V. New methods for calculating metabolic rate with special reference to protein metabolism. The Journal of Physiology. 1949;109(1–2):1–9.

93. Charpentier E, et al. 3’ RNA sequencing for robust and low-cost gene expression profiling V.1. [published online ahead of print: January 28, 2021]. https://www.protocols.io/view/3-rna-sequencing-for-robust-and-low-cost-gene-expr-n2bvj94ewlk5/v1. Accessed October 1, 2026.

94. Sadek A, et al. A conserved monocyte activation program links brain injury to systemic immune adaptation and clinical outcomes. Front Immunol. 2026;17. 10.3389/fimmu.2026.1731068.

95. Lustig Y, et al. Separation of the gluconeogenic and mitochondrial functions of PGC-1α through S6 kinase. Genes Dev. 2011;25(12):1232–1244.

96. Hardy GH. MENDELIAN PROPORTIONS IN A MIXED POPULATION. Science. 1908;28(706):49–50.

97. Weinberg W. Über vererbungsgesetze beim menschen. Zeitschrift für induktive Abstammungs-und Vererbungslehre. 1908;1(1):377–392.

