## Supplementary figures and info for "The PGC-1α Gly482Ser rs8192678 SNP reprograms oxidative metabolism to favor energy storage in females"

### 1) Supplementary Figures and Legends (1 file)

### 2) Supplementary Files (4 files)

<u>Supplementary File 1</u>*: Differential gene expression analysis between S/S and G/G mice across liver, muscle and (subcutaneous adipose tissue) SCAT.*

This file contains the results of differential expression analyses comparing S/S and G/G mice within each tissue. Reported metrics include mean normalized expression (baseMean), log2 fold change (log2FoldChange), standard error of the log2 fold change (lfcSE), Wald test statistic (stat), nominal p-value, and Benjamini-Hochberg false discovery rate (FDR)-adjusted p-value (padj). Positive log2 fold change values indicate higher expression in S/S mice relative to G/G mice, whereas negative values indicate lower expression in S/S mice. Significantly differentially expressed genes were defined as FDR-adjusted $p < 0.05$. Genes are annotated as upregulated or downregulated in S/S relative to G/G mice.

Worksheets:

- Res_df_Liver SS vs GG: Differential expression results in liver of S/S vs G/G mice.
- Res_df_Muscle SS vs GG: Differential expression results in gastrocnemius muscle of S/S vs G/G mice.
- Res_df_SCAT SS vs GG: Differential expression results in SCAT of S/S vs G/G mice.

<u>Supplementary File 2</u>. *Biological pathways commonly enriched across tissues in S/S vs G/G mice.*

Fast gene set enrichment analysis (fgsea) was performed independently in liver, gastronecmius muscle, and subcutaneous adipose tissue (SCAT) using complete gene lists ranked by DESeq2 Wald statistics from S/S versus G/G mice. This file reports biological pathways that were significantly enriched in common between tissue pairs and across all three tissues. For each pathway, the direction of enrichment in S/S relative to G/G mice is indicated as upregulated or downregulated.

Worksheets:

- Common pathway L-M-S SS vs GG: pathways significantly enriched in liver, muscle, and SCAT.
- Common pathway L-M SS vs GG: pathways significantly enriched in both liver and muscle.
- Common pathway L-SCAT SS vs GG: pathways significantly enriched in both liver and SCAT.

- Common pathway M-SCAT SS vs GG: pathways significantly enriched in both muscle and SCAT.

Supplementary File 3. *Gene set enrichment analysis results for liver, gastronecmius muscle and SCAT transcriptomes in S/S vs G/G mice.*

Fast gene set enrichment analysis (fgsea) was performed independently in liver, muscle, and subcutaneous adipose tissue (SCAT) using complete gene lists ranked by DESeq2 Wald statistics from S/S versus G/G mice. This file contains the complete pathway enrichment results for each tissue, including nominal *p*-values (pval), false discovery rate (FDR)-adjusted *p*-values (padj), enrichment scores (ES), normalized enrichment scores (NES), the number of permutations yielding enrichment scores at least as extreme as the observed score (*nMoreExtreme*), pathway size (size), and leading-edge genes (leadingEdge). Positive NES values indicate pathways upregulated in S/S mice relative to G/G mice, whereas negative NES values indicate pathways downregulated in S/S mice.

Worksheets:

- Muscle SS vs GG: complete fgsea results for gastrocnemius muscle.
- SCAT SS vs GG: complete fgsea results for subcutaneous adipose tissue.
- Liver SS vs GG: complete fgsea results for liver.

Supplementary File 4: *Metadata tables for indirect calorimetry experiments performed in male and female PGC1aSer481Gly mice.*

This file contains animal- and experiment-level metadata associated with the indirect calorimetry studies reported in this manuscript. Metadata include animal characteristics (species, genetic background, genotype, age, sex, body weight and body composition), experimental conditions (diet, fasting and HFHS refeeding protocol, ambient conditions, photoperiod, cohort information), and calorimetry acquisition parameters (equipment, acclimation period, energy expenditure calculation method, and cage enrichment).

Worksheets:

- Metadata_Males: metadata for experiments performed in male mice.
- Metadata_Females: metadata for experiments performed in female mice.

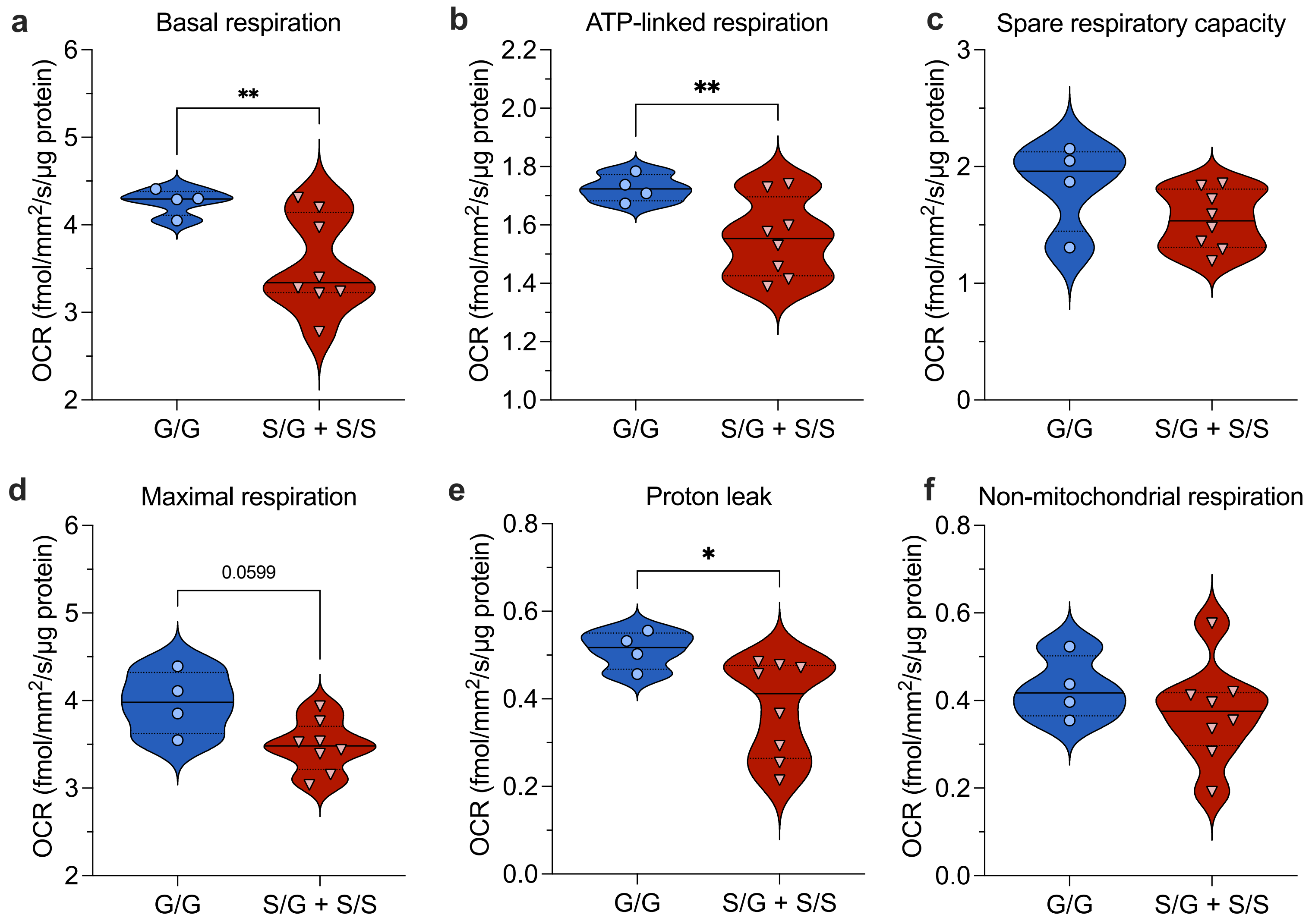


**Figure S1. The PGC-1α Gly482Ser SNP significantly reduces respiratory capacity in human iPSCs under a dominant genetic model.**

**a**) Basal respiration, **b**) ATP-linked respiration, **c**) spare respiratory capacity, **d**) maximal respiration, **e**) proton leak and **f**) non-mitochondrial respiration in G/G versus S/G + S/S human iPSCs (dominant genetic model for the serine allele, n=4 independent clones/genotype). Respiration parameters were derived from OCR measurements and normalized to total cellular protein content. Values for each clone were averaged across 3-4 independent experiments. Statistics: Unpaired two-tailed t-test with Welch's correction. $^{*}p \leq 0.05$, $^{**}p \leq 0.01$.

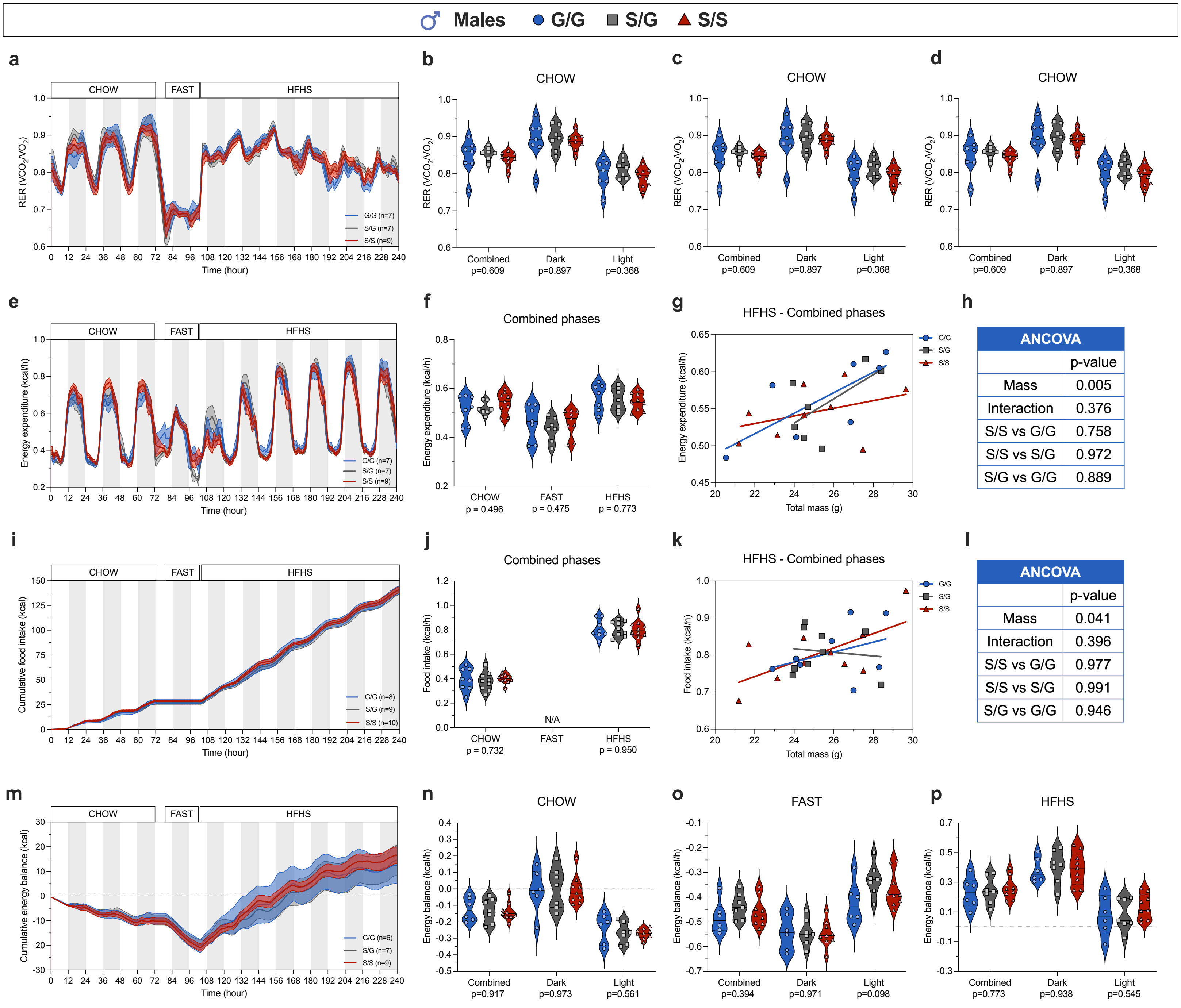


**Figure S2. Energy metabolism is similar in male PGC-1α$^{Ser481Gly}$ mice under chow, fasting and HFHS conditions.**

**a**) RER versus time in male PGC-1α$^{Ser481Gly}$ mice and violin plots representing the mean RER per mouse during **b**) chow feeding, **c**) fasting and **d**) HFHS refeeding per photoperiod. **e**) Energy expenditure (kcal/h) versus time. Violin plots representing the mean rates of **f**) energy expenditure per mouse during chow, fasting and HFHS combined photoperiods. **g**) Linear regression of mean energy expenditure rates against total body mass during the HFHS combined photoperiod and **h**) ANCOVA of energy expenditure. **i**) Cumulative food caloric intake (kcal) versus time. Violin plots representing the mean rates of **j**) food intake (kcal/h) per mouse during chow, fasting, and HFHS combined photoperiods. **k**) Linear regression of mean food intake rates against total body mass during the HFHS combined photoperiod and **l**) ANCOVA of food intake. **m**) Cumulative energy balance (kcal) versus time. Violin plots representing the mean energy balance rates (kcal/h) per mouse during **n**) chow, **o**) fasting, and **p**) HFHS feeding per photoperiod. Shaded areas represent dark phases. Statistics: Genotype effects were assessed using **b,c, n-p**) one-way ANOVA, **d**) Kruskal-Wallis test, or **f,j**) ANCOVA with total body mass as a covariate, performed independently for each photoperiod or nutritional state.

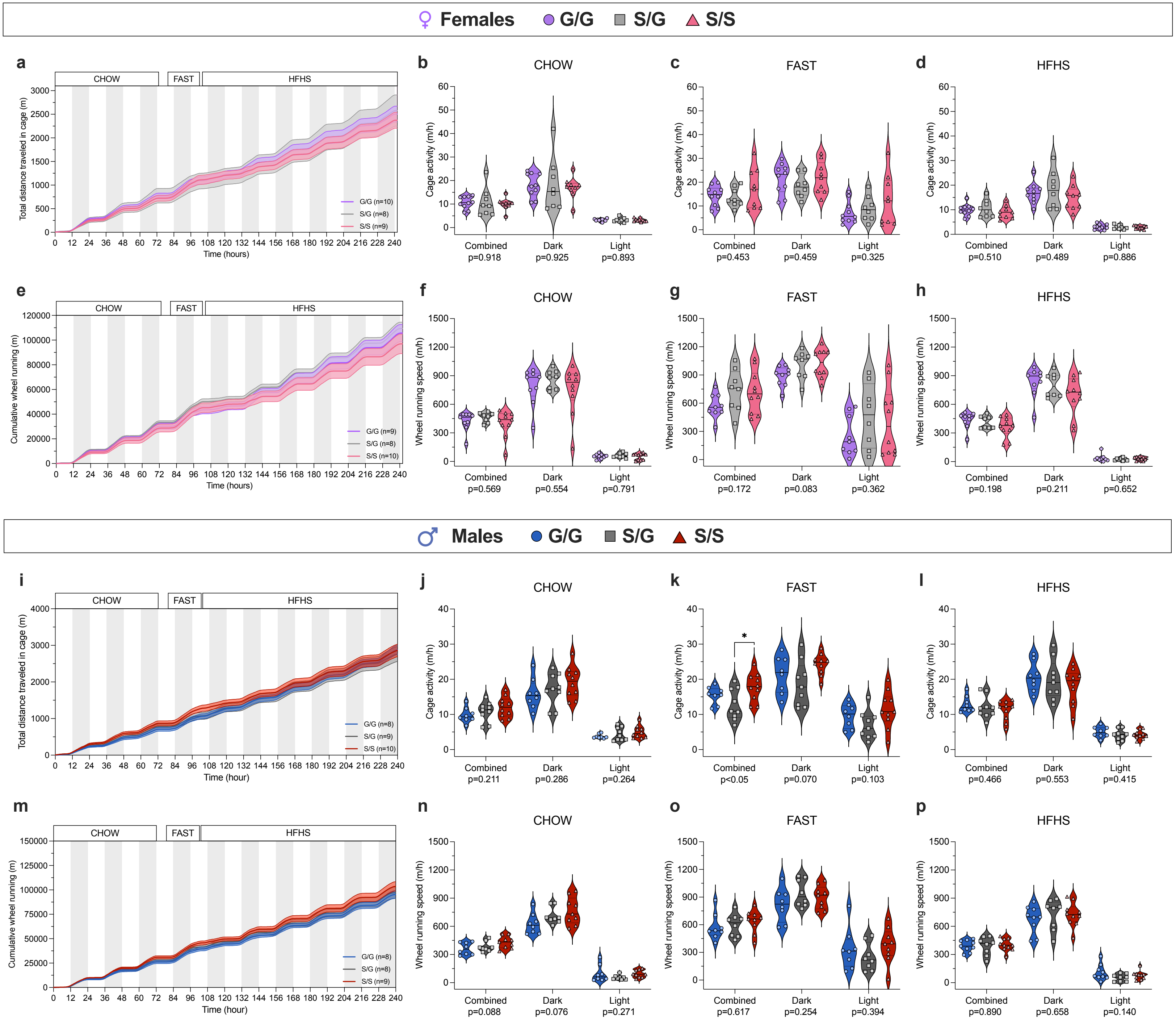


**Figure S3. Voluntary physical activity is increased in male and female S/S mice during fasting.** Total distance traveled in the cage (m) versus time in **a**) female and **i**) male PGC-1α$^{Ser481Gly}$ mice. Violin plots representing the average cage activity (m/h) per mouse during **b,i**) chow feeding, **c,k**) fasting, and **d,l**) HFHS feeding in female and male mice, respectively. Cumulative distance run on the wheel (m) versus time in **e**) female and **m**) male mice. Average wheel running speed (m/h) during **f,n**) chow feeding, **g,o**) fasting, and **h,p**) HFHS feeding in female and male mice, respectively. Shaded areas represent dark phases. Statistics: Genotype effects were assessed using **b-d, g, j-l, n-p**) one-way ANOVA with Tukey's post-hoc test or **f,h**) Kruskal-Wallis test, performed independently per photoperiod. *$p \leq 0.05$.

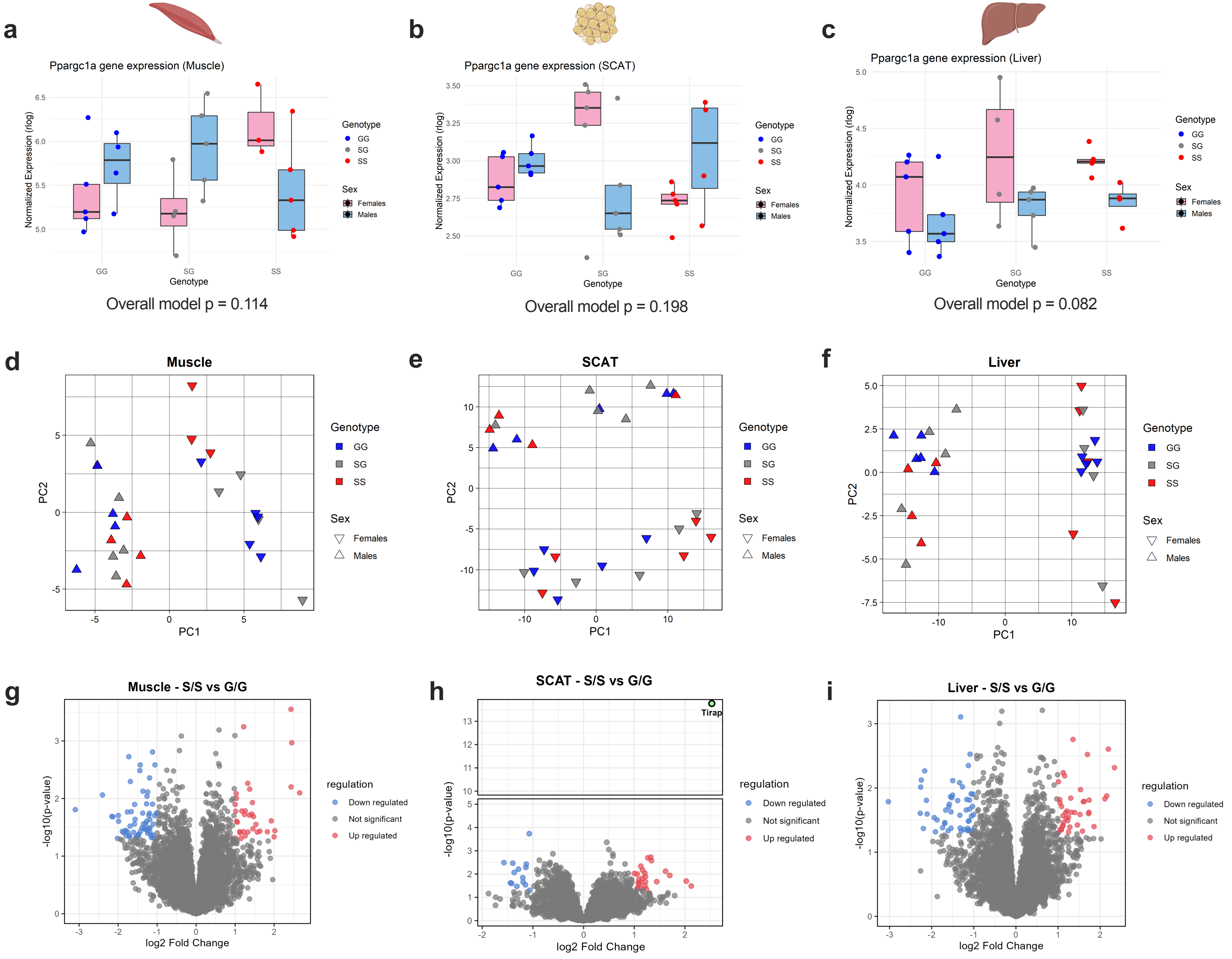


**Figure S4. Sex is the major driver of transcriptional variation across muscle, SCAT and liver tissues from PGC-1α$^{Ser481Gly}$ mice.**

**a-c**) *Ppargc1α* rlog-transformed normalized expression values in each tissue from male and female PGC-1α$^{Ser481Gly}$ mice, stratified by sex and genotype. Principal component analysis (PCA) of transcriptome profiles from male and female PGC-1α$^{Ser481Gly}$ mice in **d**) muscle (PC1 = 32% expl.var, PC2 = 18% expl.var), **e**) SCAT (PC1 = 42% expl.var, PC2 = 18% expl.var), and **f**) liver (PC1 = 75% expl.var, PC2 = 4% expl.var). Volcano plot showing the distribution of differentially expressed genes (DEGs) in S/S vs G/G mice in **g**) muscle, **h**) SCAT and **i**) liver. Upregulated genes are shown in red, downregulated genes in blue, and non-significant genes in grey. DEGs were defined as $|log_2FC| > 1$ and raw p-value ≤ 0.05 by DESeq2. n = 9-10 G/G, 9-10 S/G, 8-9 S/S per tissue.

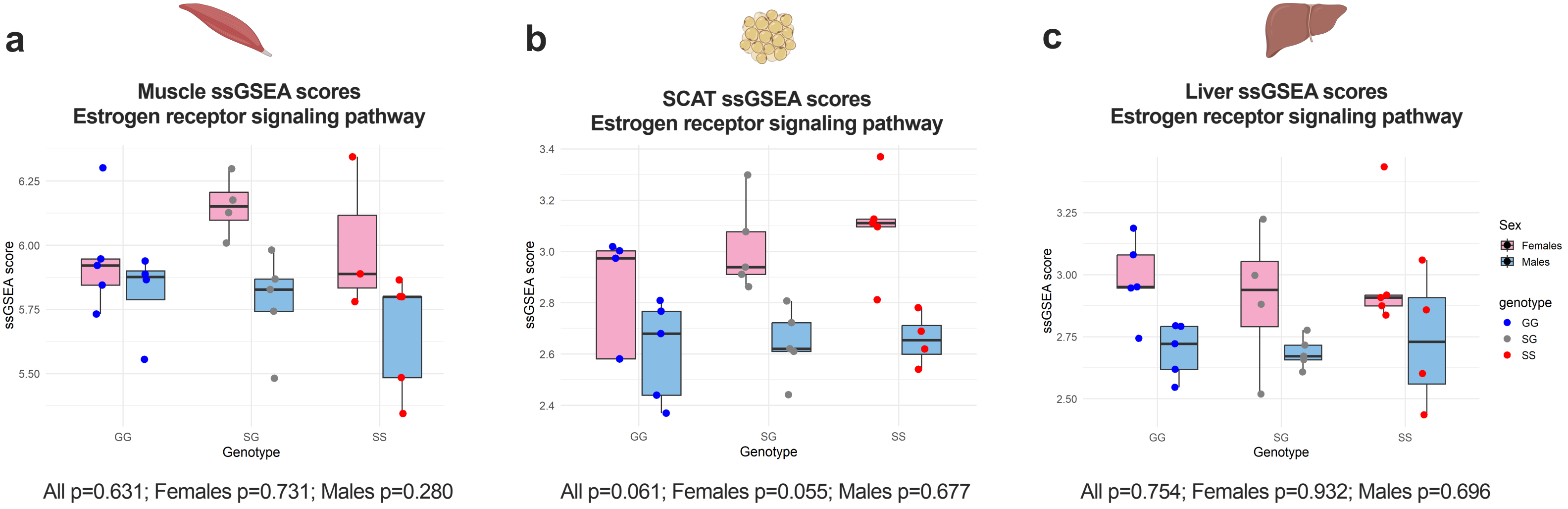


**Figure S5. Estrogen receptor signaling pathways tends to be higher in SCAT of S/S females.**

ssGSEA scores for the estrogen receptor signaling pathway in **a**) muscle, **b**) SCAT, and **c**) liver of PGC-1α$^{Ser481Gly}$ mice in both sexes. Each point represents an individual animal; boxplots indicate ssGSEA scores across genotype, and statistical significance were evaluated using limma moderated t-test (eBayes) and reported using FDR adjusted p-value. n = 9-10 G/G, 9-10 S/G, 8-9 S/S per tissue.
